# Rab11 endosomes coordinate centrosome number and movement following mitotic exit

**DOI:** 10.1101/2021.08.11.455966

**Authors:** Nikhila Krishnan, Maxx Swoger, Michael Bates, Judy Freshour, Peter J. Fioramonti, Alison Patteson, Heidi Hehnly

**Affiliations:** Syracuse University, Department of Biology, 107 College Place, Syracuse, NY 13244, USA; Syracuse University, Department of Physics, Syracuse, Physics Building, Syracuse, NY 13244, USA

**Author notes:** Lead contact, correspondence, Twitter: @LovelessRadio.

## Abstract

The last stage of cell division involves two daughter cells remaining interconnected by a cytokinetic bridge that is cleaved in a process called abscission. During pre-abscission, we identified that the centrosome moves in a Rab11-dependent manner towards the cytokinetic bridge in human cells grown in culture and in an *in vivo* vertebrate model, *Danio rerio* (zebrafish). Rab11-endosomes are dynamically organized in a Rab11-GTP dependent manner at the centrosome during pre-abscission and this organization is required for the centrosome protein, pericentrin, to be enriched at the centrosome. Using zebrafish embryos, we found that reduction in pericentrin expression or optogenetically disrupting Rab11-endosome function inhibited centrosome movement towards the cytokinetic bridge and abscission resulting in daughter cells prone to being binucleated and/or having supernumerary centrosomes. These studies suggest that Rab11-endosomes contribute to centrosome function during pre-abscission by regulating pericentrin organization resulting in appropriate centrosome movement towards the cytokinetic bridge and subsequent abscission.

## INTRODUCTION

During the onset of cell division, the centrosome duplicates to build a bipolar mitotic spindle, which is a macromolecular machine that ensures each daughter cell receives an equal complement of chromosomes. Following chromosome separation, cytokinetic furrow formation, and the dismantling of the bipolar spindle, the daughter cell centrosomes were noted to migrate from one end of the cell towards the cytokinetic bridge. This movement in a human cell line (HeLa) is required for cytokinetic bridge cleavage, referred to as abscission (Piel et al., 2001). A separate study demonstrated in a variety of different mammalian cell lines that centrioles have a range of described motions with some directed toward the bridge and other motions being highly irregular (Jonsdottir et al., 2010). Our studies expand upon these previous findings, where we have identified a potential mechanism involving the small GTPase Rab11 in centrosome motility towards the cytokinetic bridge during pre-abscission and that this mechanism is conserved *in vivo* using the vertebrate model, *Danio rerio* (zebrafish).

The recycling endosome (RE) and its associated small GTPase, Rab11, localize to a specific sub-structure of the centrosome, mother centriole appendages and this association was shown to be required for RE function (Hehnly et al., 2012; Naslavsky and Caplan, 2020). Strikingly, if RE function was disrupted, centrosome function was also disrupted resulting in a loss of microtubule populations at metaphase centrosomes (Hehnly and Doxsey, 2014) or cilia formation during G0 (Xie et al., 2019). Based on this relationship between Rab11-REs and the centrosome during these cell cycle stages, we hypothesize that the centrosome requires an association with Rab11-REs during pre- abscission for their orientation towards the cytokinetic bridge. Centrosome movement during pre-abscission is similar to the movement observed by the centrosome as cells undergo duplicated centrosome migration during *Drosophila melanogaster* neuroblast divisions (Lerit and Rusan, 2013). During neuroblast divisions, the centrosome duplicates during S-phase where one centrosome migrates to the distal side of the cell to mature. This migratory process requires a pericentrin-like protein (PLP) (Lerit and Rusan, 2013). Our studies herein identified that centrosome reorientation towards the cytokinetic bridge uses mechanisms involving both pericentrin and Rab11-endsosomes.

Rab11 is required for successful abscission. After furrow ingression, dividing animal cells stay interconnected for some time by a narrow intercellular bridge that contains a proteinaceous structure known as the midbody (reviewed in (Chen et al., 2012)). Rab11-associated membranes were identified to transport with their associated cargo into the cytokinetic bridge. These vesicles can fuse and potentially prime the membranes next to the midbody for an abscission event (Schiel et al., 2012). When depleting Rab11 using siRNAs (Wilson et al., 2005) or inhibiting the ability of Rab11- associated vesicles to transport into the bridge using optogenetics (Rathbun et al., 2020a), abscission failure occurs both in cell culture and in the zebrafish embryo. However, the relationship between Rab11-associated endosomes and the centrosome during this process has not been investigated.

## RESULTS AND DISCUSSION

### Mitotic centrosomes containing Rab11-endosomes reorient towards the cytokinetic bridge during pre-abscission

We found that REs associate dynamically with the centrosome as it moves towards the cytokinetic bridge. To determine this, we first tested whether centrosome reorientation towards the cytokinetic bridge is a conserved process. Dividing zebrafish embryonic cells expressing two different centrosome markers, centrin-GFP (Rathbun et al., 2020b; Zolessi et al., 2006) or PLK1-mCherry (Rathbun et al., 2020a), were imaged (Figure 1A, S1A, Video S1) and compared to human (HeLa) cells expressing the centrosome markers DsRed-PACT (a c-terminal centrosome targeting domain taken from pericentrin, Figure 1B, S1B, Video S2) or centrin-GFP (Figure 1C, (Kuo et al., 2011; Piel et al., 2001)). In 74.7% of cases one of two centrosomes reoriented towards the cytokinetic bridge in zebrafish embryonic cells (Figure 1A, 1C, Video S1), compared to 86.18% in human cells (Figure 1B, 1C, Video S2). We then examined the spatial and temporal association of REs with the centrosome during pre-abscission. The Rab11-effector protein FIP3 was tagged with GFP (Wilson et al., 2005) and expressed in HeLa cells with a centrosome marker, DsRed-PACT (Figure 1B). We found a population of REs at the centrosomes during anaphase and this population significantly increased as the cell progressed through cytokinesis and pre-abscission (Figures 1B, S1C, S1E, Video S2). This same population of Rab11-REs was noted at the centrosome in pre-abscising zebrafish embryo cells (Figure 1D) and this organization was similar to what was reported for another RE associated GTPase, Rab25 (Willoughby et al., 2021). These findings suggest REs organization at the centrosome and centrosome movement towards the cytokinetic bridge is a conserved process during pre-abscission.

**Figure 1:**
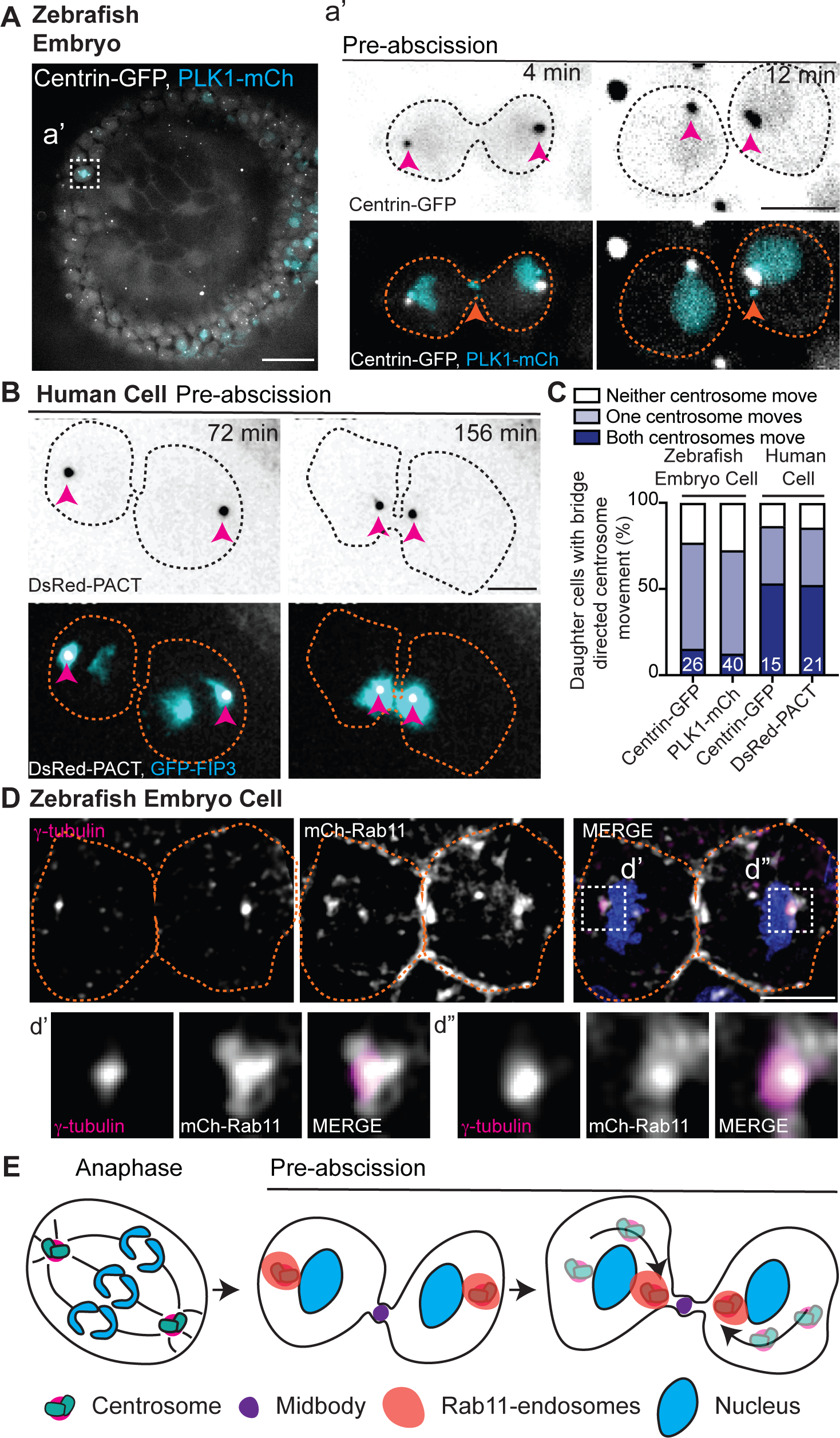
Mitotic centrosomes containing Rab11-endosomes reorient towards the cytokinetic bridge during pre-abscission. (A) Zebrafish embryo with centrin-GFP (gray) and PLK1-mCherry (cyan). Scale bar, 50 μm. (a’) Inset of metaphase cell in A. Time-lapse of centrin-GFP (inverted grays, top panel; grays, bottom) and PLK1-mCherry (cyan). Video S1. (B) Time-lapse of pre- abscising human (HeLa) cell expressing DsRed-PACT (inverted grays, top panel; gray, bottom) and GFP-FIP3 (cyan, bottom). Video S2. (A, B) Scale bar, 10 μm. (A-B) Pink arrow, centrosome. Orange arrow, midbody. Dashed lines, cell boundaries. (C) Percentage of daughter cells with centrosome movement towards the cytokinetic bridge of a zebrafish embryo (3 hpf) and human (HeLa) cells. n-values, Table S1. (D) A zebrafish embryo pre-abscising cell expressing mCherry-Rab11 (gray) fixed and immunolabeled for γ-tubulin (magenta) and DNA (DAPI, blue). Dashed lines, cell boundaries. Scale bar, 10 μm. Insets (d’ and d’’), 2X magnification. (E) Model depicting centrosomes (green) containing Rab11-endosomes (orange) reorienting towards the cytokinetic bridge with associated midbody (purple) during pre-abscission.

Human cells were used to understand the temporal and spatial distribution pattern of REs due to their optical accessibility. Using live cell imaging we identified a second population of REs that form adjacent to the cytokinetic bridge following the formation of the centrosome population, the centrosome then reorients with a population of REs to this second RE population (Figures 1B, 1E, S1C, Video S2). Another membrane organelle, the Golgi apparatus (labeled with MannII-Ruby), is dispersed in small puncta during cytokinesis and starts to form two separate fragmented compartments next to the centrosome and the bridge (Figure S1D). This localization pattern is different from the REs that remain tightly organized at the centrosome and adjacent to the bridge (Figure S1C-D). To test if the second population of REs originated from the centrosome population of REs, a photo-convertible Dendra-Rab11 was used. The centrosome population of Dendra-Rab11 endosomes were photoconverted from a 507 nm emission to a 573 nm emission by placing a region of interest (ROI) over the centrosome where 405 nm light was applied. A significant population of 573 nm REs formed the second population of REs adjacent to the bridge (Figure S1F-G), suggesting that the second population of REs originated from the centrosome population of REs.

### Rab11 GTPase function and centrosome localization is required for centrosome bridge-directed movement

To assess the robustness of centrosome reorientation during pre-abscission across scales (cell culture to zebrafish embryos), we developed a way to normalize distance traveled by the centrosome in relation to the cell (Figure S2A). This involved recording the movement of the centrosome and the cell body as vectors and using vector subtraction to calculate centrosome movement within the reference frame of the cell in both cell culture (Figure S2B) and in zebrafish embryo cells (Figure S2C). Using this, we analyzed distance traveled by the centrosome during pre-abscission and whether Rab11 is required. Since Rab11 and its associated endosomes have been reported to directly associate with the centrosome (Hehnly et al., 2012) and can contribute to mitotic centrosome structure (Hehnly and Doxsey, 2014), we examined if Rab11 is required for centrosome reorientation by first using CRISPR to create a Rab11-null centrin-GFP or GFP-FIP3 HeLa cell line (Figure S2D). Similar to previous reports, cells null for Rab11 resulted in increased binucleated cells indicative of an abscission failure (Rathbun et al., 2020a; Wilson et al., 2005) that was rescued with ectopic expression of mCherry-Rab11 (Figure S2E, S2F).

In control GFP-FIP3 expressing pre-abscising cells, endogenous Rab11 colocalizes with its effector protein, FIP3. In Rab11-null cells, no Rab11 is detected by immunohistochemistry and GFP-FIP3 is no longer organized in a tight centrosome localized compartment (Figure 2A). Rab11-null cells were less likely to move at least one of their centrin-GFP decorated centrosomes towards the cytokinetic bridge (51.43% of cells) compared to control cells (80%, Figure 2B, 2C, Video S3). When measuring the total distance traveled by centrosomes during pre-abscission, we found that centrosomes traveled a significantly shorter distance in Rab11-null cells (32.19 ± 2.03 μm), compared to control (46.50 ± 2.01 μm), Figure S2G). From this movement, we identified that the centrosome had significantly more cytokinetic bridge directed movement (distance traveled 8.64 ± 1.63 μm) in control cells compared to Rab11-null cells (2.00 ± 0.86 μm, Figure 2F), suggesting that Rab11 is required for directed centrosome movement towards the cytokinetic bridge.

**Figure 2:**
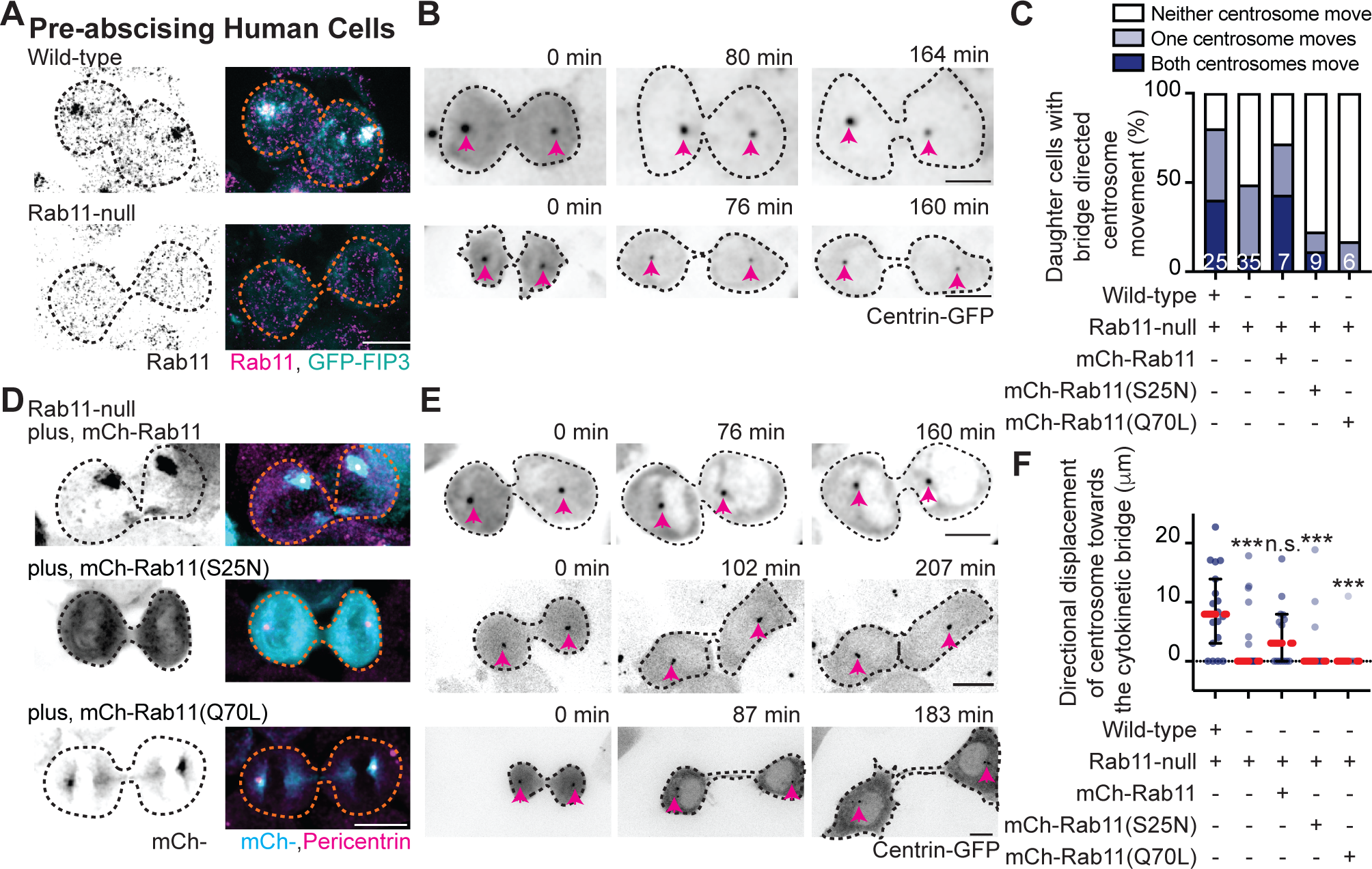
Rab11 GTPase function and centrosome localization is required for centrosome bridge directed movement. (A) Wild-type and Rab11-null GFP-FIP3 (cyan, merge) human (HeLa) cells fixed and immunostained for Rab11 (inverted grays, left; magenta, merge). (B) Time-lapse of wild- type and Rab11-null centrin-GFP (inverted grays) pre-abscising human (HeLa) cells. (A- Pink arrows, centrosome (B). Dashed lines, cell boundaries (A-B). Scale bar, 10 μm. Video S3. (C-F) Rab11-null cells expressing mCherry-Rab11, -Rab11 (S25N) or - Rab11(Q70L). (C) Percentage of cells with cytokinetic bridge directed centrosome (centrin-GFP) movement (stacked bar graph). (D) GFP-FIP3 Rab11-null cells expressing mCherry-Rab11, -Rab11 (S25N) or -Rab11(Q70L) (inverted grays, left; cyan, merge) fixed and immunostained for pericentrin (magenta, merge). (E) Centrin-GFP (inverted grays) Rab11-null pre-abscising human (HeLa) cells expressing mCherry-Rab11, - Rab11(S25N) or -Rab11(Q70L). (D-E) Pink arrows, centrosome (E). Dashed lines, cell boundaries (D-E). Scale bar, 10 μm. Video S4. (F) Distance traveled by centrosome towards cytokinetic bridge (scatter plot). Median (orange dashed line) and quartiles (dark lines) shown. One-way ANOVA with Dunnett’s multiple comparison to wildtype, ***p<.001. (C, F) Statistical results detailed in Table S2.

To test the requirement for Rab11 GTPase function, we expressed fluorescent tagged versions of Rab11 in Rab11-null cells that include: wild-type Rab11 (mCherry- Rab11), a mutant that mimic’s the GDP-bound state of Rab11 (mCherry-Rab11(S25N)), or a mutant that mimics the GTP-bound state of Rab11 (mCherry-Rab11(Q70L), Figure 2D). Expression of mCherry-Rab11, -Rab11(S25N), and -Rab11(Q70L) were all comparable to endogenous Rab11 expression in control cells (Figure S2D). mCherry- Rab11 and -Rab11(Q70L) had similar centrosome localization patterns compared to - Rab11(S25N) that remained more cytosolic (Figure 2D). Expressing mCherry-Rab11 in Rab11-null cells rescued centrosome bridge directed movement in 71.43% of cells with a bridge directed distance traveled of 4.57 ± 1.46 μm, compared to -Rab11(S25N) (22.22% of cells, directed distance of 1.93 ± 1.17 μm) or -Rab11(Q70L) (16.67% of cells, directed distance of 0.92 ± 0.92 μm) (Figure 2C, 2E, 2F, Video S4). We were surprised that mCherry-Rab11(Q70L) did not partially rescue centrosome movement towards the cytokinetic bridge since it can mimic the active state of Rab11 and localizes to centrosomes. One possibility is that GTP to GDP cycling is required for Rab11 to regulate centrosome reorientation and affects the temporal and spatial organization of Rab11 during pre-abscission. To test this, we examined whether a difference in mCherry-Rab11 and -Rab11(Q70L) dynamics occurred using Fluorescent Recovery After Photobleaching (FRAP) by photobleaching the population of mCherry-Rab11 or -Rab11(Q70L) at the centrosome and comparing the fluorescent recovery over time. We found significant differences in dynamics between mCherry-Rab11 and -Rab11(Q70L) (Figure S2H-S2K). Rab11(Q70L) cells had a decreased mobile fraction of 36.57 ± 4.48%, compared to Rab11 at 59.02 ± 5.08%. Rab11(Q70L) also presented with a decreased half-life (2.27 ± 0.33s) compared to Rab11 (9.00 ± 1.31s, Figure S2I-S2K). Collectively, these findings suggest that mCherry-Rab11 dynamics at centrosomes is partly regulated by its ability to cycle GTP to GDP contributing to centrosome movement towards the cytokinetic bridge.

### Rab11 GTP-cycling is required for the centrosome protein, pericentrin, centrosome localization

We find that γ-tubulin and pericentrin partially localize to REs during pre- abscission in both human and zebrafish embryo cells (Figure 3A-B), which is consistent with previous studies that identified pericentrin and γ-tubulin at REs during metaphase in human cells (Hehnly and Doxsey, 2014). We specifically find that REs associate with pericentrin and γ-tubulin at and outside of the centrosome (Figure 3A-B, inset demonstrating acentrosomal localization). In human cells expressing the RE marker, GFP-FIP3, and the centrosome targeting domain from pericentrin, DsRed-PACT, GFP- FIP3 colocalized with acentrosomal DsRed-PACT. These GFP-FIP3/DsRed-PACT puncta move towards and combine with the main centrosome (Figure 3C) suggesting that REs may help contribute to centrosome overall organization and function.

**Figure 3:**
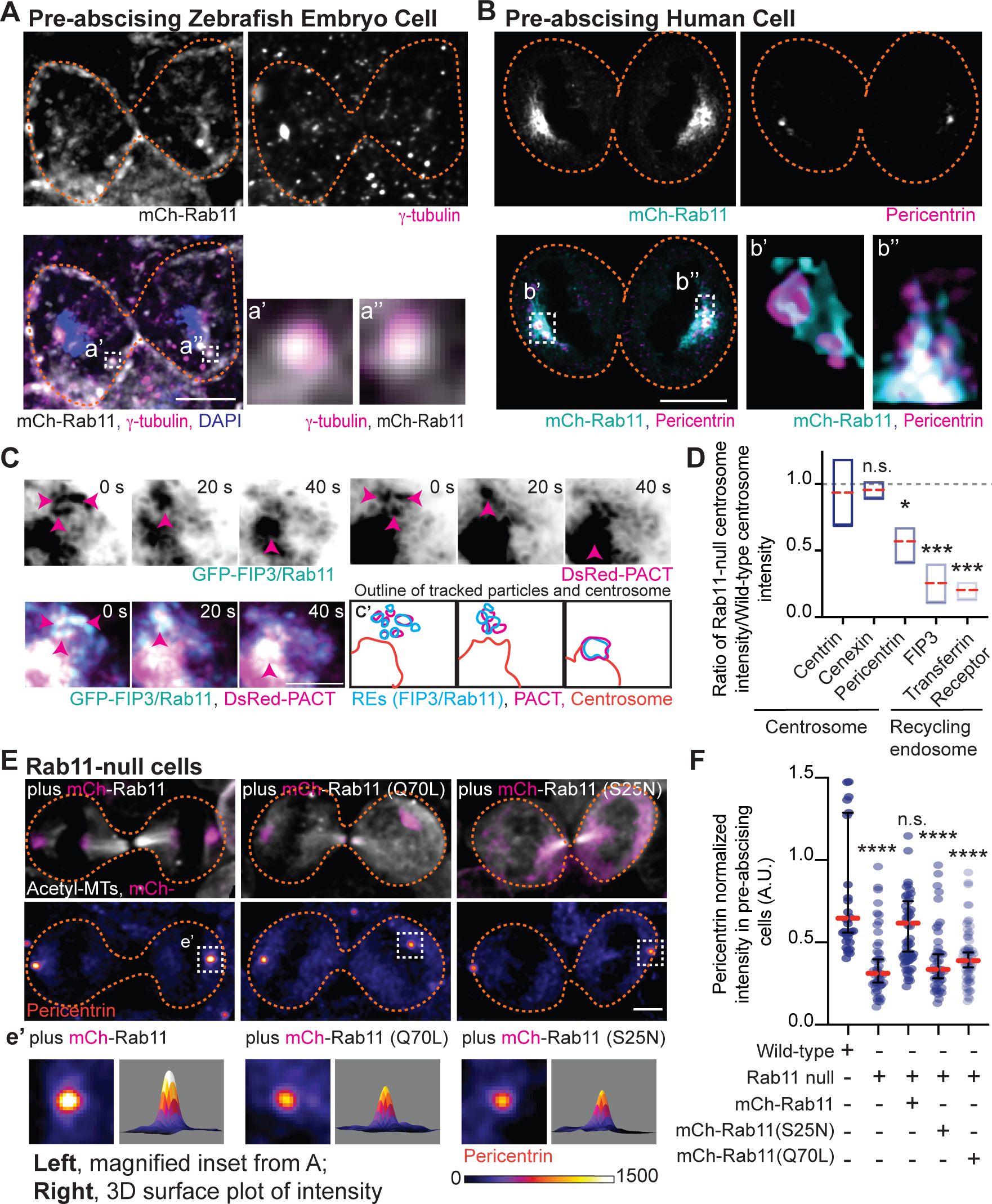
Rab11-GTP cycling is required for centrosome protein, pericentrin, centrosome localization. (A-B) Pre-abscising zebrafish embryonic cell (A) or a Rab11-null human (HeLa) cell expressing mCherry-Rab11 (gray, A and B top; cyan, B) fixed and immunostained for γ- tubulin (magenta, A), pericentrin (magenta, B), and DNA (DAPI, blue, A). Scale bar, 10 μm. Magnified insets of acentrosomal site (a’, a”, b”) and centrosome (b’). (C) Time-lapse of GFP-FIP3/Rab11 (inverted grays, left; cyan, merge) centrosome region in a pre- abscising human (HeLa) cell expressing DsRed-PACT (inverted grays, left; magenta, merge). Pink arrows, acentrosomal fragments positive for GFP-FIP3/Rab11/DsRed- PACT. Scale bar, 10 μm. (c’) Outline of tracked particles noted by pink arrow. (D) Box and whisker plot with mean (orange dashed line) depicting ratio of Rab11-null centrosome intensity over wild-type centrosome intensity of cenexin, centrin, pericentrin, FIP3, and transferrin receptor. 25^th^ and 75^th^ percentiles noted by boxed boundaries. One-way ANOVA with Dunnett’s multiple comparison to centrin, n.s. not significant, *p<0.05 and ***p<0.001. (E) GFP-FIP3 Rab11-null pre-abscising human (HeLa) cells ectopically expressing mCherry-Rab11, -Rab11(S25N) or -Rab11(Q70L) (magenta, top) were fixed and immunolabelled for acetylated-tubulin (gray, top) and pericentrin (fire LUT, bottom). Scale bar, 10 μm. (e’) 3x magnified centrosome inset (left), 3D fluorescent intensity surface plot of inset (right). (F) Scatter plot with median (orange dashed line) and quartiles (dark lines) depicting normalized pericentrin intensities at centrosomes from pre- abscising human (HeLa) cells. One-way ANOVA with Dunnett’s multiple comparison to wild-type control, n.s. not significant, ****p<0.0001. (D, F) Statistical results detailed in Table S3.

Since Rab11-null pre-abscising cells are unable to orient their centrosome towards the cytokinetic bridge, we analyzed whether localization of centrosome components and RE components at the centrosome was altered in Rab11-null cells compared to control cells (Figures 3D-3F, S3A-C). Fluorescent intensity was measured in fixed cells using a ROI placed over the centrosome and compared between control and Rab11-null cells (as in (Hehnly et al., 2012)). The fluorescent values were plotted as a ratio of Rab11-null over control cells (Figure 3D). A ratio of 1 implies no difference between Rab11-null cells and wild-type, but a ratio significantly less than 1 suggests decreased centrosome localization in Rab11-null cells. Rab11-null cells had significantly decreased centrosome localized GFP-FIP3 (Figures 3D, S3A, S3B), the RE cargo, transferrin receptor (Figures 3D, S3B), and pericentrin (Figures 3D, S3A). However, the centriole appendage protein, cenexin, and centriole protein, centrin, were not affected by Rab11 loss (Figures 3D, S3C), suggesting that Rab11-REs are involved in PCM maintenance but not centriole organization. Pericentrin intensity at the centrosome was rescued in Rab11-null cells by expression of mCherry-Rab11, but not mCherry-Rab11(S25N) or -Rab11(Q70L) (Figure 3E, 3F). Pericentrin expression levels were unaltered in Rab11-null cells, or cells rescued with mCherry-Rab11, -Rab11(S25N), or -Rab11(Q70L) (Figure S2D), suggesting that the targeted localization of pericentrin is disrupted with loss of Rab11 function. Taken together, Rab11 GTPase function is required for centrosome reorientation during pre- abscission (Figure 2) by facilitating pericentrin organization at the centrosome (Figure 3).

### Pericentrin and Rab11 endosomes coordinate centrosome movement and number during mitotic exit

Since loss of Rab11 caused a decrease in pericentrin levels at the centrosome in human cells (Figure 3), we tested the role of pericentrin and Rab11- endosomes in regulating centrosome movement during pre-abscission *in vivo*. To do this we used heterozygous pericentrin (*pcnt*^+/-^) zebrafish embryos (Sepulveda et al., 2018) positive for the centrosome marker centrin-GFP (embryo generation modeled in Figure S4A) or by acutely inhibiting Rab11-associated vesicles through an optogenetic oligomerization approach that relies on a hetero-interaction between CRY2 and CIB1 that is induced by the application of blue light (experiment design modeled in Figure S4B, (Rathbun et al., 2020a). *Pcnt*^+/-^ embryos were used due to our inability to obtain reliable *pcnt* ^null^ embryos. When comparing *pcnt*^+/-^ and Rab11 optogenetically clustered embryos to control embryos (non-injected or CRY2 injected) at 3.3 hours post fertilization (hpf), we identified that *pcnt*^+/-^ and Rab11 clustered embryos were significantly impaired at reorienting their centrosomes towards the cytokinetic bridge during pre-abscission (18.75% reoriented for *pcnt*^+/-^ and 26.32% for Rab11-clustered) compared to control conditions (76.91% for non-injected, and 52.37% for CRY2 injected, Figure 4A-4C, confirmation of genotype in Figure S4C, CRY2 injected control in Figure S4D). *Pcnt*^+/-^ and Rab11 clustered embryos also demonstrated significant defects in centrosome movement during pre-abscission, where the calculated distance traveled across 12 minutes was on average between 10-12 μm for *pcnt*^+/-^ and Rab11 clustered embryos compared to 15 μm with control conditions (Figure 4D). These studies suggest that both pericentrin and Rab11 are required for centrosome bridge directed motility.

**Figure 4:**
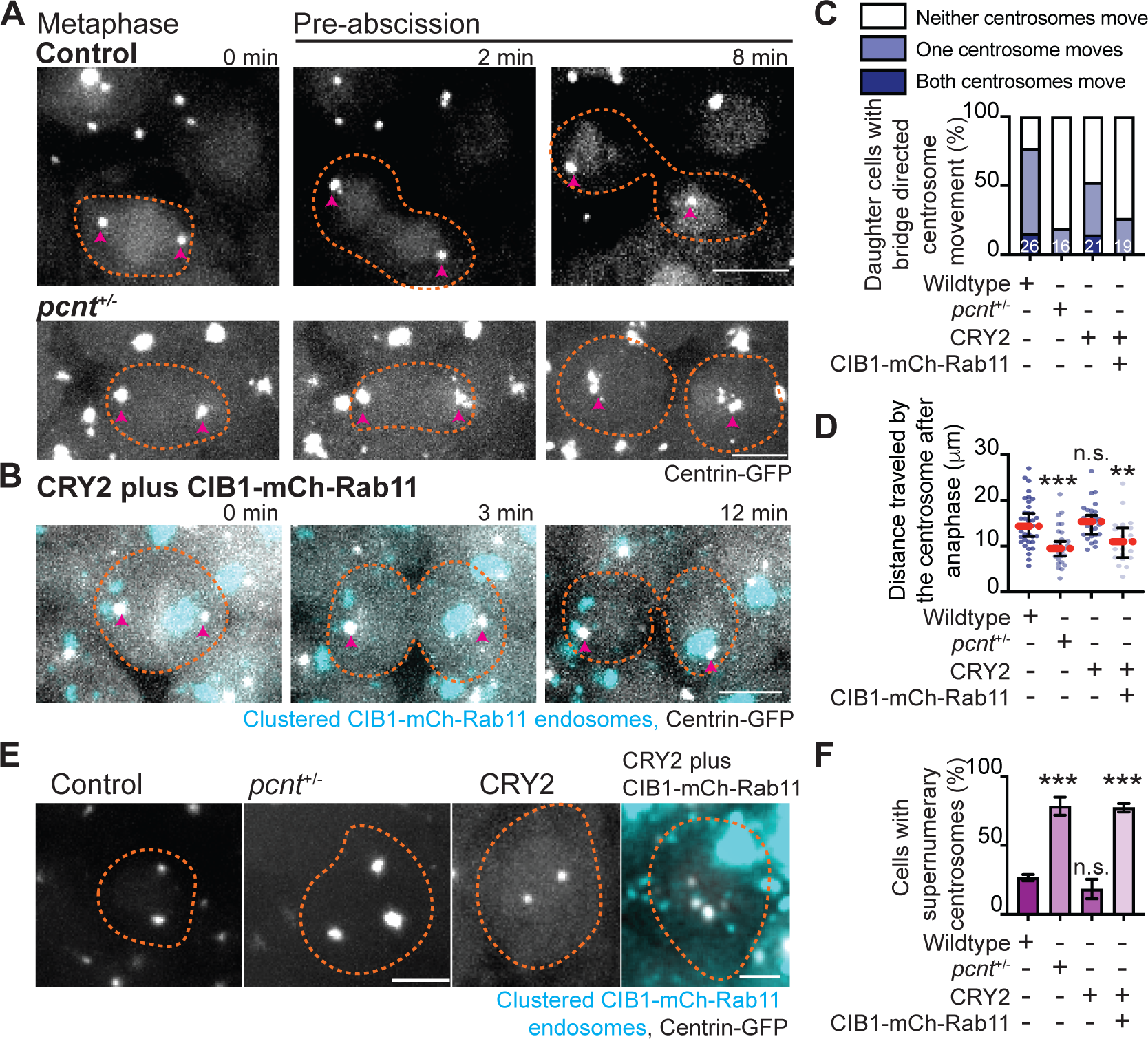
Pericentrin and Rab11 coordinate centrosome movement and number during mitotic exit *in vivo*. (A-B) Time-lapse imaging of control cells (A), *pcnt*^+/-^ cells (A), and cells with clustered Rab11 endosomes (B, cyan) from a centrin-GFP (gray) embryo. Pink arrows, centrosome. Dashed orange line, cell boundaries. Scale bar, 10 μm. (C-D) Percentage of daughter cells with cytokinetic bridge directed centrosome movement (stacked bar graph, and distance traveled of centrosome towards the cytokinetic bridge (scatter plot with median, orange dashed line, and quartiles, dark lines, D) in wildtype, *pcnt*^+/-^, CRY2 injected, or CRY2 plus CIB1-Rab11 injected embryos. (D) One-way ANOVA with Dunnett’s multiple comparison, n.s. not significant, **p<0.01, and ***p<0.001. (E-F) Interphase centrin-GFP (gray) zebrafish embryos with n=2 centrosomes or supernumerary (n>2 centrosomes) in wildtype, *pcnt*^+/-^, CRY2 injected, CRY2 plus CIB1- Rab11 injected embryos (cyan, CIB1-mCherry-Rab11 plus CRY2). (E) Orange dashed lines, cell boundaries. Scale bar, 5μm. (F) Percentage of cells with supernumerary centrosomes (n>2 centrosomes). Two-tailed student’s t-test, ***p<0.001. (C, D, F) Statistical results detailed in Table S4.

We next examined whether decreasing pericentrin (*pcnt*^+/-^) and acutely clustering Rab11-vesicles resulted in abscission defects. When cells are unable to complete abscission, cells become binucleated (Carter, 1967) or can present with supernumerary centrosomes (Pihan et al., 2003). Most cells contain either one or two centrosomes, depending on their cell cycle stage. A G1 cell contains a single centrosome composed of two centrioles. During S phase, two new (daughter) centrioles assemble near the pre- existing (mother) centriole creating two centrosomes that will move apart to create a bipolar mitotic spindle. When cells fail abscission, the two daughter cells combine gaining an extra centrosome and present as binucleated. Having too many centrosomes can result in multipolar divisions, chromosome segregation defects, defects in asymmetric cell division, loss in cell polarity, induction of invasive protrusions, and inappropriate activation of signaling pathways (Godinho and Pellman, 2014). In our studies, we counted embryonic cells that had an abnormal number of centrosomes (3 or more) or were binucleated. With *pcnt*^+/-^ embryos, we find a significant increase in embryos with binucleated cells (Figure S4E, S4G) and supernumerary centrosomes (78.33% ± 6.509 of cells) compared to control (26.57%± 2.293, Figure 4E-4F). With optogenetic clustering of Rab11-REs we find a similar dramatic increase in both the percentage of cells containing supernumerary centrosomes (77.18%± 3.029) compared to CRY2 injected controls (18.41% ± 7.079, Figure 4E-F) and binucleated cells (Figure S4F, S4G), suggesting that Rab11-endosomes and pericentrin are required for abscission.

Taken together, these studies present a scenario where Rab11-endosomes regulate centrosome function by maintaining appropriate pericentrin levels at the centrosome. When pericentrin levels are decreased, either through the loss of Rab11 function (Figure 3) or through removing one allele of *pcnt* (Figure 4), we find that cells in pre-abscission no longer reorient their centrosomes towards the cytokinetic bridge and ultimately fail abscission resulting in cells containing supernumerary centrosomes.

## ACKNOWLEDGEMENTS

We thank Li-En Jao lab (UCSD) for the Tg(*pcnt*^tup2^) transgenic zebrafish lines and genotyping protocols for characterization. This work was supported by National Institutes of Health grants R01GM127621 (H.H.) and R01GM130874 (H.H.). We thank the Blatt BioImaging Center for the use of the LSM 980 with Airyscan2 (NIH S10 OD026946-01A1). This work was also supported by the U.S Army Medical Research Acquisition Activity through the FY16 Prostate Cancer Research Programs under Award no. W81XWH-20- 1-0585 (H.H.). Opinions, interpretations, conclusions, and recommendations are those of the authors and not necessarily endorsed by the Department of Defense.

## AUTHOR CONTRIBUTIONS

N.K., H.H. designed, performed and analyzed experiments, wrote manuscript; H.H oversaw project and edited manuscript; A.P., M.S. edited manuscript and advised on centrosome tracking; M.B., P.J.F., J.F. performed experiments and associated analyses.

## STAR METHODS

### Resource Availability

#### Lead contact

For further information or to request resources/reagents, contact the Lead Contact, Heidi Hehnly

*Materials availability:* New materials generated for this study are available for distribution.

*Data and Code Availability:* All data sets analyzed for this study are displayed.

### Experimental Model and Subject Details

*Fish Lines:* Zebrafish lines were maintained using standard procedures approved by Syracuse IACUC (Institutional Animal Care Committee) (Protocol #18-006). Embryos were raised at 28.5°C and staged (as described in ref. (Kimmel et al., 1995)). Wildtype and/or transgenic zebrafish lines used for live imaging and immunohistochemistry are listed in key resource table.

### Method Details

#### Antibodies

Antibody catalog information used in HeLa cells and zebrafish embryos are detailed in key resource table.

#### Plasmids and mRNA

Plasmids were generated using Gibson Cloning methods (NEBuilder HiFi DNA assembly Cloning Kit) and maxi-prepped before injection and/or transfection. mRNA was made using mMESSAGE mMACHINE^TM^SP6 transcription kit. See key resource table for a list of plasmid constructs and mRNA used.

#### Cell Culture

HeLa cells stably expressing GFP-FIP3 or centrin-GFP (from (Hehnly and Doxsey, 2014; Hehnly et al., 2012; Kuo et al., 2011; Piel et al., 2001; Wilson et al., 2005)) were maintained at 37°C with 5% CO2. Rab11A CRISPR vector and Rab11A HDR vector were transfected into cells using Mirus TransIT-LT1 transfection reagent (key resource table). Cells were selected in puromycin (5 μg/ml). Rab11-null cells were transfected with mCherry-Rab11 (pCS2-mCherry-Rab11), dominant negative Rab11 (pCS2-mCherry- Rab11(S25N)) and constituently active Rab11 (pCS2-mCherry-Rab11(Q70L)) using Mirus TransIT-LT1. Cells were tested for Rab11 levels using a Western blot.

#### Western blot

Cell lysates were acquired by suspending cells in lysis buffer (HSEG buffer pH 7.4: 40mM HEPES, 40mM NaCl, 5mM EDTA, 4% Glycerol, 20mM NaF; 1% TritonX- 100; 1X protease inhibitor; 0.1mM PMSF). After collecting post-nuclear supernatant from lysates, protein concentration was calculated using Bio-Rad Protein Assay Kit II (see key resource table). Standard Western blot procedures were performed. Nitrocellulose membranes were probed with primary antibody and/or primary antibody conjugated to horseradish peroxidase diluted in TBS-Tween20 and incubated overnight at 4°C. The membranes were probed using appropriate secondary antibody for an hour at room temperature. The protein levels were visualized using Clarity^TM^ Western ECL substrate (see key resource table) and imaged using Bio-Rad ChemiDoc^TM^ imager.

#### Immunofluorescence

Cells were plated on #1.5 coverslips until they reach 90% confluence fixed in 4% PFA at room temperature (30 min) or 100% ice cold methanol (10 mins). Standard immunofluorescent procedures were performed for PFA fixation (Hehnly et al., 2006) and for methanol (Colicino et al., 2018). Cover slips were rinsed with dH2O and mounted on glass slides using either Prolong Diamond with DAPI mounting media or Prolong Gold (see key resource table). For zebrafish embryo immunofluorescent protocols see (Aljiboury et al., 2021).

#### Genotyping pcnt^+/-^ zebrafish

Tail fins of adult zebrafish were clipped, whole embryos or fixed and stained embryos were used to extract genomic DNA and genotyped according to (Sepulveda et al., 2018).

#### Imaging

Zebrafish and tissue culture cells were imaged using Leica DMi8 (Leica, Bannockburn, IL) equipped with a X-light V2 Confocal unit spinning disk equipped with a Visitron VisiFRAP-DC photokinetics unit, a Leica SP8 (Leica, Bannockburn, IL) laser scanning confocal microscope (LSCM) and/or a Zeiss LSM 980 (Carl Zeiss, Germany) with Airyscan 2 confocal microscope. The Leica DMi8 is equipped with a Lumencore SPECTRA X (Lumencore, Beaverton, OR), Photometrics Prime-95B sCMOS Camera, and 89 North-LDI laser launch. VisiView software was used to acquire images. Optics used with this unit are HC PL APO x40/1.10W CORR CS2 0.65 water immersion objective, HC PL APO x40/0.95 NA CORR dry and an HCX PL APO x63/1.40-0.06 NA oil objective. The SP8 laser scanning confocal microscope is equipped with HC PL APO 20x/0.75 IMM CORR CS2 objective, HC PL APO 40x/1.10 W CORR CS2 0.65 water objective and HC PL APO x63/1.3 Glyc CORR CS2 glycerol objective. LAS-X software. was used to acquire images. The Zeiss LSM 980 is equipped with a T-PMT, GaAsP detector, MA-PMT, Airyscan 2 multiplex with 4Y and 8Y. Optics used with this unit are PL APO 63X/1.4 NA oil DIC. Zeiss Zen 3.2 was used to acquire the images. A Leica M165 FC stereomicroscope equipped with DFC 9000 GT sCMOS camera was used for staging and phenotypic analysis of zebrafish embryos.

For live human cell imaging, cells were plated on #1.5 glass bottom Ibidi slides or MatTek plates (see key resource table). Cells were imaged using spinning disk confocal, LSCM, or wide-field fluorescent microscopy followed by deconvolution (AutoQuant X3). Cells were imaged in a temperature and CO2 controlled chamber for 1-10 hours at 0.5 - 4-minute time intervals.

For zebrafish embryo imaging, fluorescent transgenic or mRNA injected embryos (refer to strains and mRNAs in key resource table, and for injection protocols refer to (Aljiboury et al., 2021)) were embedded in 2% agarose at 3.3-5 hours post fertilization (hpf) and imaged using the spinning disk or LSCM.

#### Fluorescence Recovery after photobleaching (FRAP) and photoconversion

FRAP experiments were conducted 24 hours post transfection of mCherry-Rab11 or - Rab11(Q70L) in centrin-GFP cells using the Leica DMi8 with spinning disk and photokinetics unit (Visitron VisiFRAP-DC). A region of interest (ROI) was marked at the centrosome in a cytokinetic cell and a 405nm laser was used to photobleach mCherry within that region. Following photobleaching the cell was imaged live to identify recovery of fluorescent signal at the centrosome at 3 second intervals for 3 minutes. The ImageJ FRAP calculator macro plug-in was used to generate FRAP curves and calculate half-life and immobile fraction values. Graphs were generated using PRISM9 software.

For photoconversion experiments Dendra-Rab11 was expressed in HeLa cells. A ROI was placed over centrosome localized Dendra-Rab11 in a single daughter cell during pre-abscission. A 405 nm laser was applied within the ROI to photoconvert Dendra- Rab11 from green emission (507 nm) to a red emission (573 nm).

#### Rab11 optogenetics experiments in zebrafish

Tg (-5actb2:cent4-GFP), Tg (sox17:GFP-CAAX), Tg *BAC*(cftr-GFP) zebrafish embryos were injected with 50-100pg of CIB1- mCherry-Rab11, CIB1-mCerulean-Rab11, CRY2-mCherry and/or CRY2 mRNA at the one cell to 4 cell stage (Rathbun et al., 2020a). Embryos were allowed to develop to a minimum of 3.5 hpf and exposed to 488nm light while being imaged using the spinning disk confocal microscope.

#### Tracking centrosome movement

Human (HeLa) cells and zebrafish embryos expressing centrin-GFP were imaged using widefield or confocal based imaging. Cells were projected and the manual tracking plug-in (FIJI/Image J) was used to track the movement of the centrosome (centrin-GFP) from metaphase exit until 3 hours post anaphase in HeLa cells and up to 12 mins post anaphase or abscission completion in zebrafish embryos. The X and Y coordinates of the centrosome and cell body were recorded at each time-point. The change in position of X and Y of the cell body, marked as the center of the cell at each time point), (ΔX_Cellbody_ = X_cellbody-t2_-X_Cellbody-t1_; ΔY_Cellbody_ = Y_Cellbody-t2_-Y_Cellbody- t1_)) was subtracted from the change in position of X and Y of the centrosome (ΔX_Centrosome_ = X_Centrosome-t2_-X_Centrosome-t1_; ΔY_Centrosome_ = Y_Centrosome-t2_-Y_Centrosome-t1_) between each time-point to control for the movement of the centrosome resulting from the motion of the cell body (ΔX = ΔC_Xellbody_-ΔX_Centrosome;_ ΔY = ΔY_Cellbody_-ΔY_Centrosome_). Using Pythagorean theorem, net distance was calculated between time-points 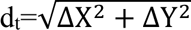, which were then added together to calculate the total distance travelled by the centrosome 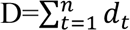; where n=final time point. For directional distance towards the cytokinetic bridge, centrin-GFP positive centrosomes were tracked from when they reach the polar ends of the daughter cells to when they reach the cytokinetic bridge using the method mentioned above. If the centrosome does not move towards the cytokinetic bridge then the distance traveled by that centrosome is recorded as 0 μm.

#### Centrosome Intensity profiles

Z stacks shown are maximum projected representative cells (FIJI/ImageJ). For intensity calculations, z-stacks were sum projected, a ROI was marked around the centrosome, and mean fluorescence intensities were measured. Fluorescent intensities were calculated as mean intensity – minimum intensity. Intensities calculated were then normalized to average intensity of the parent cell population within the experiment. Three-dimensional intensity profiles were created using FIJI/ImageJ. Outliers were identified using an iterative Grubb’s test with alpha = 0.05 using PRISM9 software.

#### Phenotypic analysis of cells exhibiting cytokinetic defects

Human cells and zebrafish embryo cells (described above) were assessed for the presence of binucleated cells, represented as a percentage.

#### Tracking centrosome number

Zebrafish embryo cells (described above) were assessed for abnormalities in the number of centrosomes at interphase. The number of centrosomes within each cell was counted and the percentage of cells with greater than two centrosomes were graphed using PRISM9 software.

#### Statistical Analysis

Unpaired, two-tailed t-tests and one way ANOVA were performed using PRISM9 software. **** denotes a p-value< 0.0001, *** p-value<0.001, ** p- value<0.01, * p-value<0.05. For further information on detailed statistical analysis see supplemental tables S1, S2, S3, S4.

## KEY RESOURCE TABLE

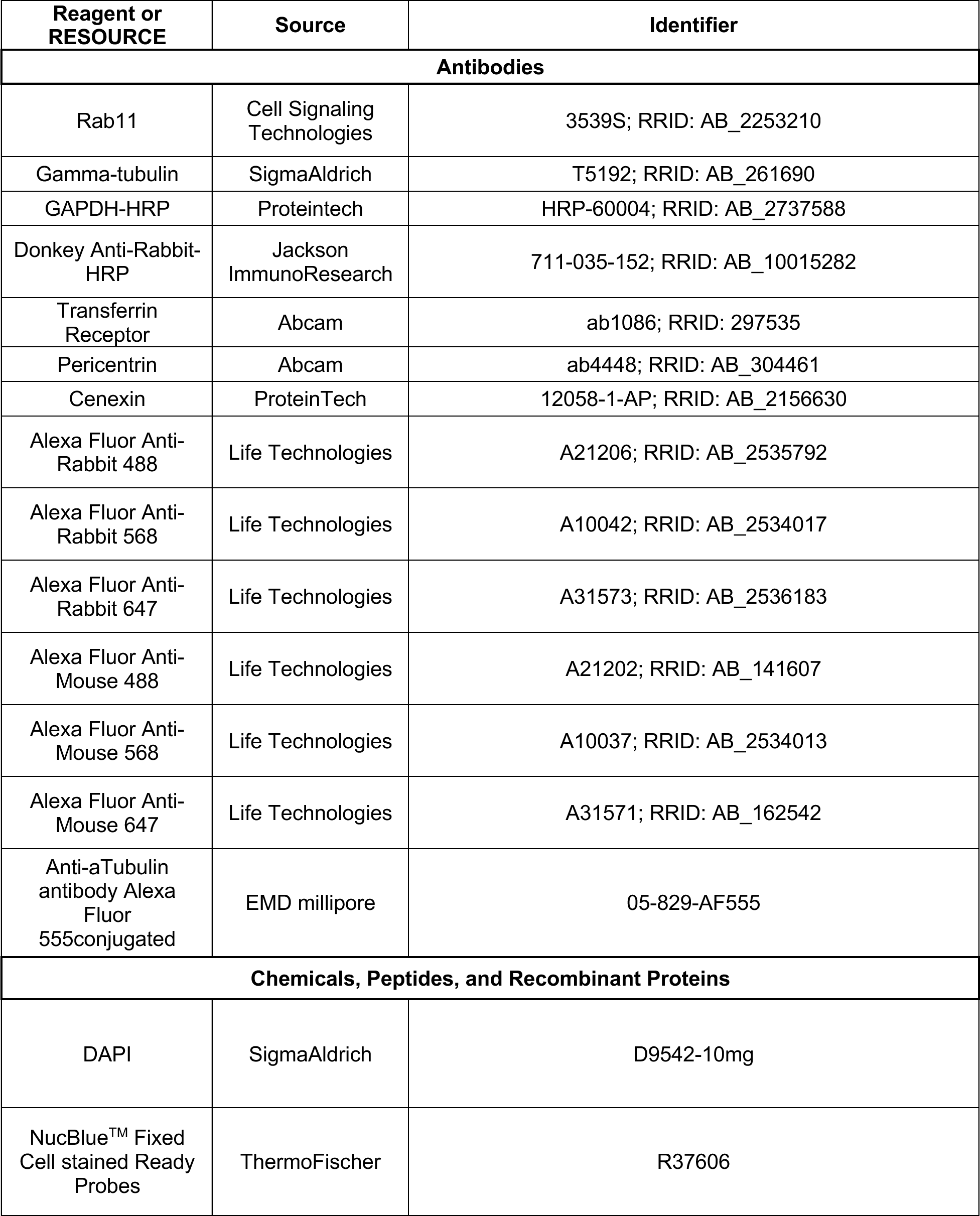
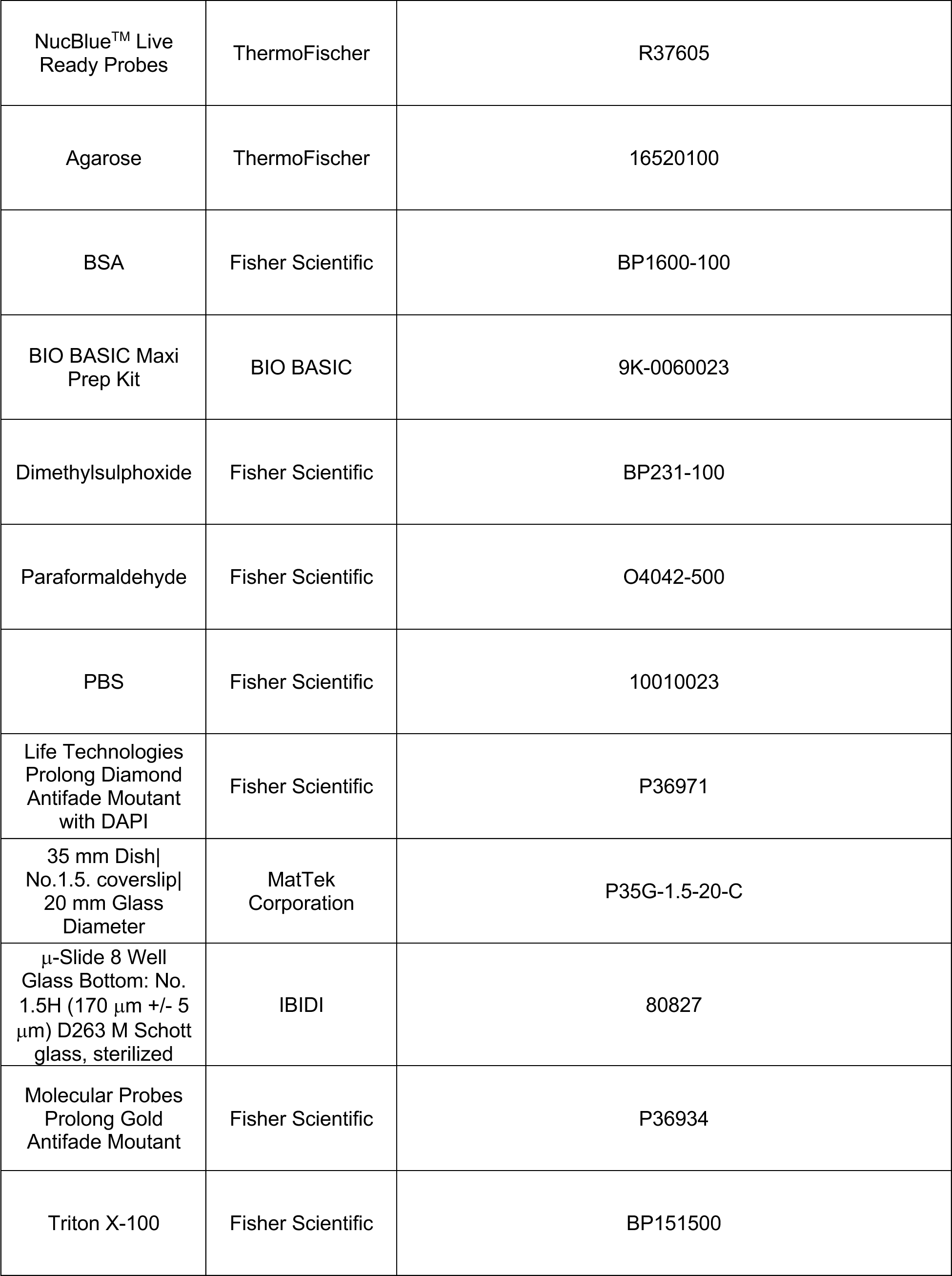
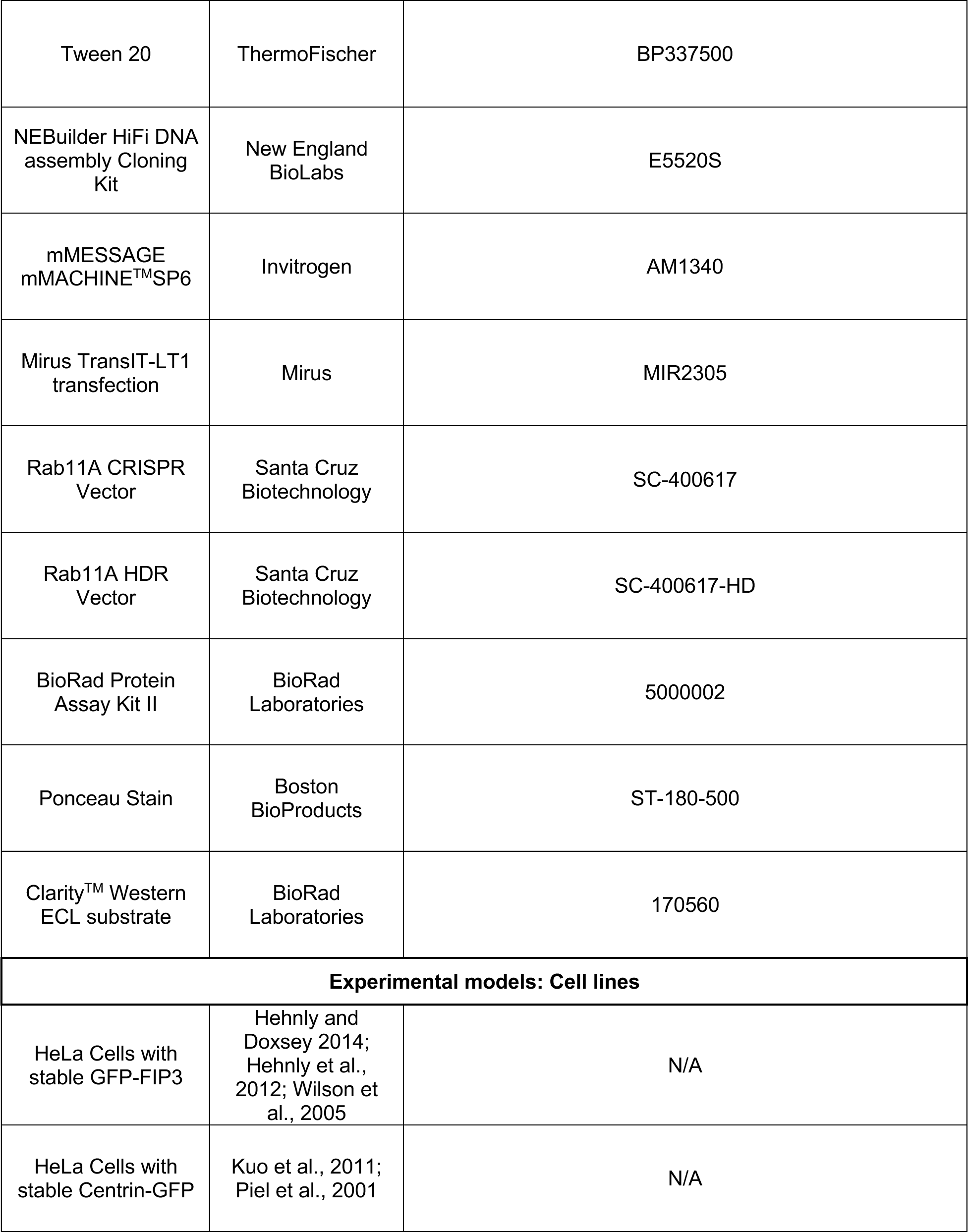
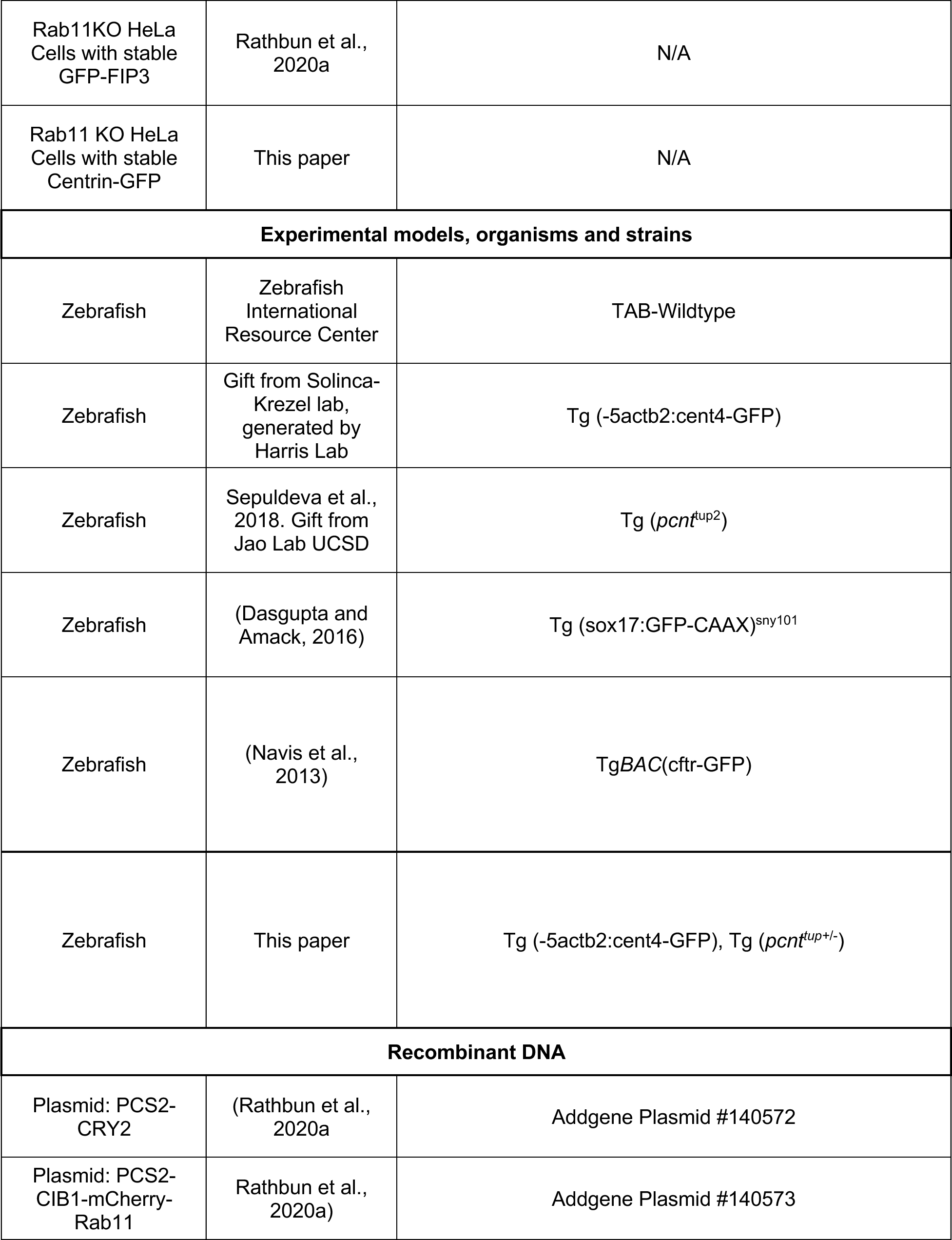
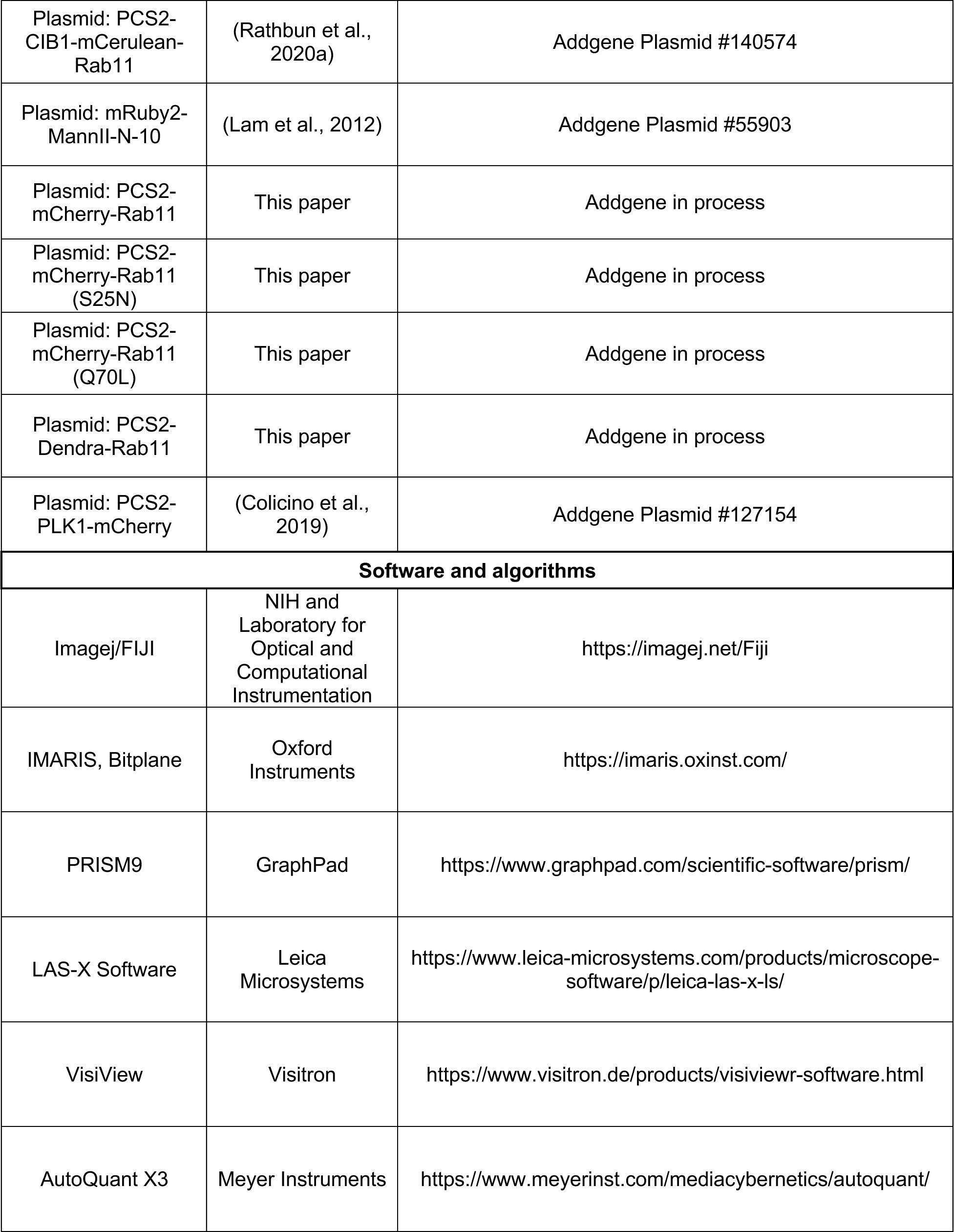
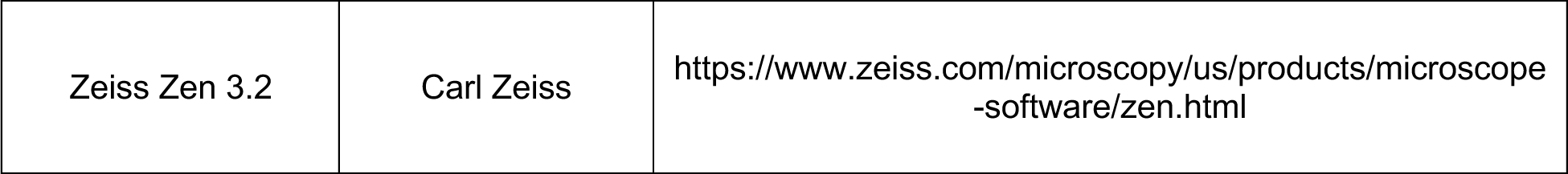

---

**Figure S1:**
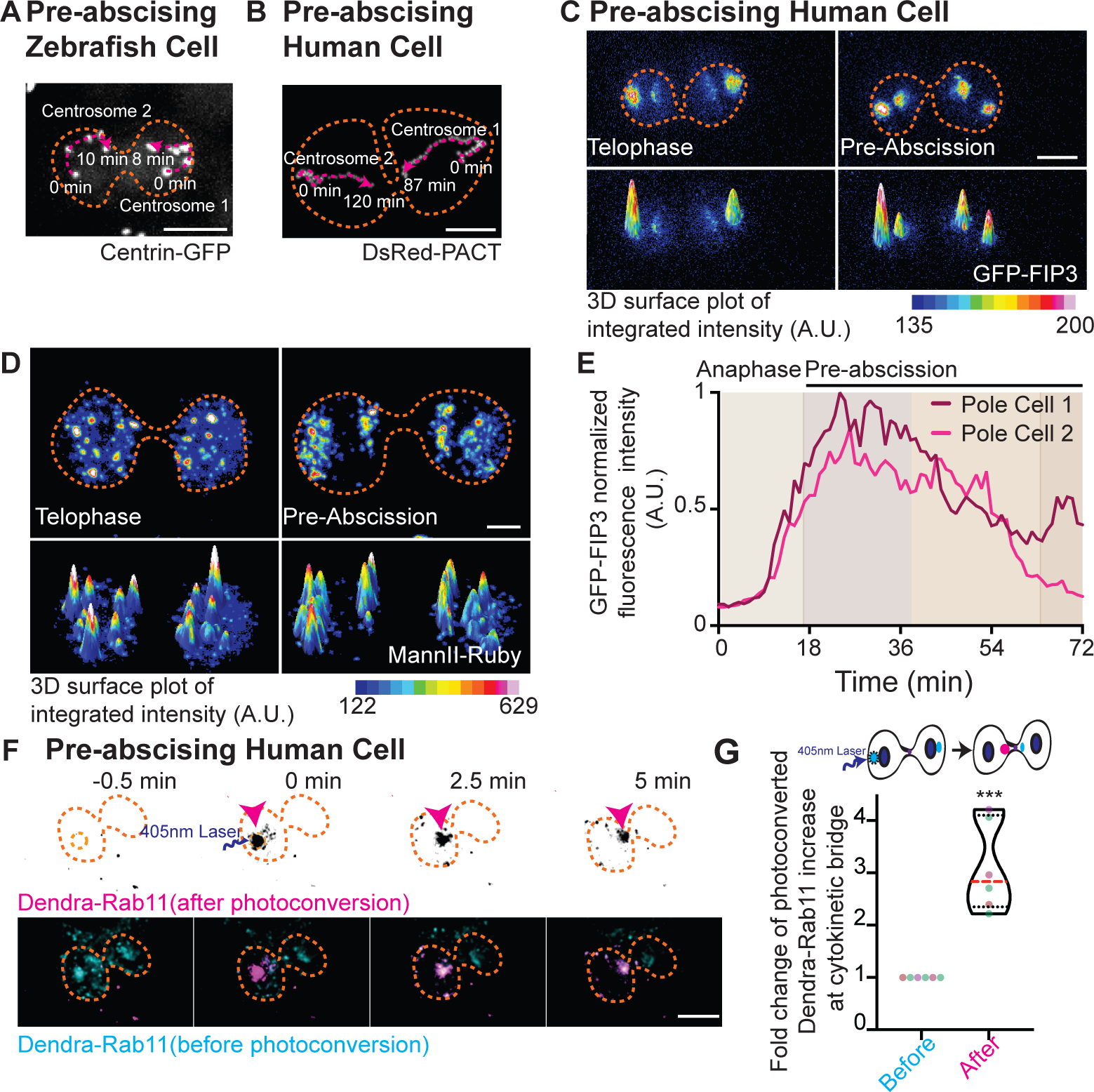
Mitotic centrosomes containing Rab11-endosomes reorient towards the cytokinetic bridge during pre-abscission. (A-B) Overlay of centrosome movement within pre-abscising zebrafish embryo cell (A) and human (HeLa) cell (B) over time from Figure 1A and 1B. The centrosome is labeled with centrin-GFP (magenta), orange dashed lines, cell boundaries. Scale bar, 10 μm. (C- Time-lapse of a pre-abscising human (HeLa) cell expressing GFP-FIP3 (16-color LUT, C) or MannII-Ruby (16-color LUT, D). Bottom panels depict three-dimensional surface plot of the intensities portrayed in top panels. Dashed lines, cell boundaries. Scale bar, 5 μm. (E) Line graph depicting GFP-FIP3 normalized fluorescence intensity from anaphase exit to late abscission at the polar compartments from (C). Intensities of the brighter pole (pole 1), dark magenta and the less bright pole (pole 2), light pink. (F) Time-lapse of a pre-abscising human (Hela) cell expressing Dendra-Rab11. Dendra-Rab11 is photoconverted from 507 nm emission (cyan, merge) to 573 nm emission (inverted grays, top row; magenta, merge) using 405nm light within an ROI over the centrosome (pink arrow) at 0 min. Dashed lines, cell boundaries. Scale bar, 10 μm. (G) Violin plot with median (orange dashed line) and quartiles (black lines) depicting fold change of photoconverted Dendra-Rab11 (507 nm emission) increase at the cytokinetic bridge. Each dot in the plot represents a cell and the different colors depict separate experiments. n=6 cells and n=4 experiments. Two-tailed student’s t-test, ***p<0.001. Statistical results detailed in Table S1.

**Figure S2:**
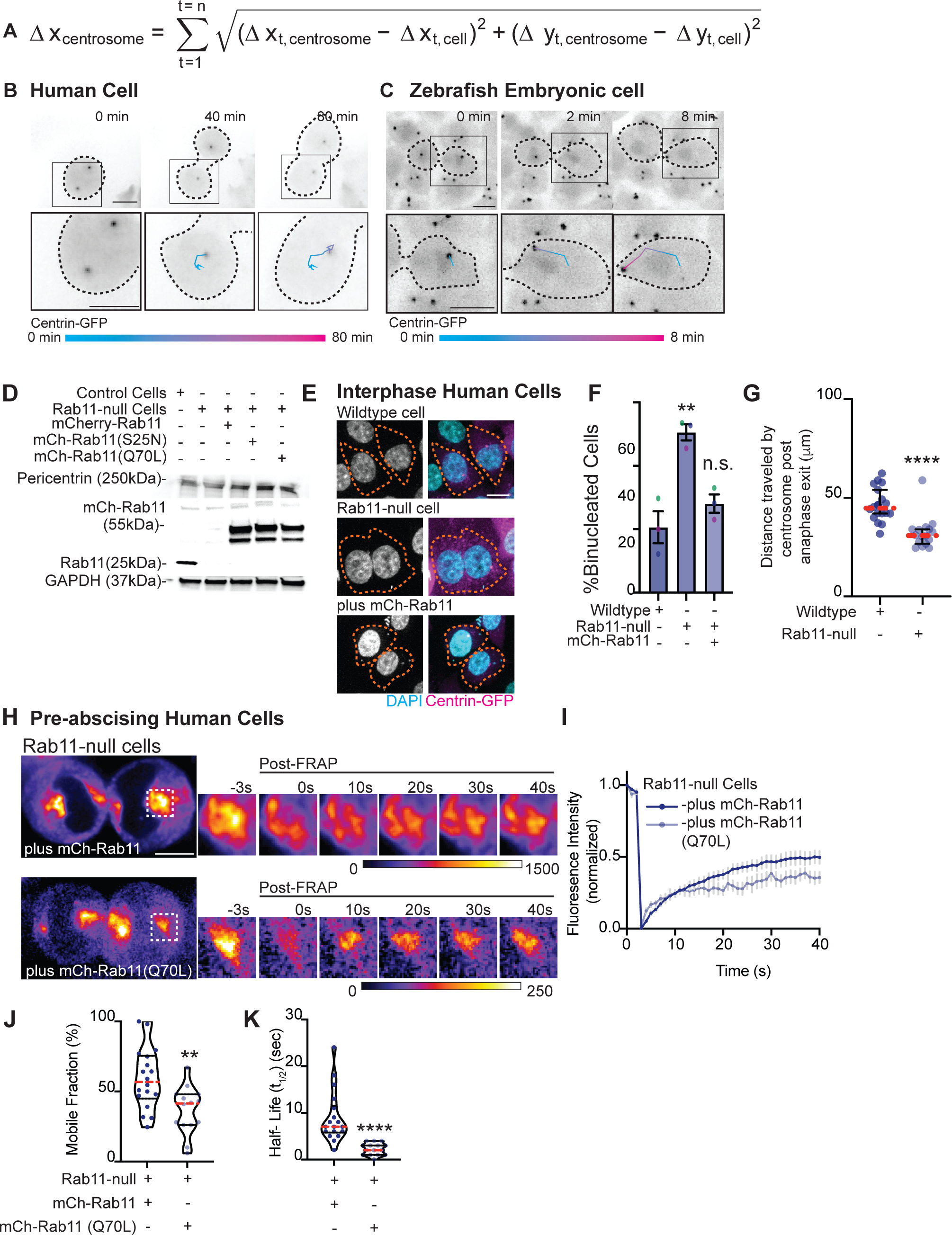
Rab11 GTPase function and centrosome localization is required for centrosome bridge directed movement. (A) Equation to calculate distance traveled by centrosome in relation to the cell. This involves recording movement of the centrosome and the cell body as vectors and using vector subtraction to calculate centrosome movement within the reference frame of the cell to accurately calculate distance traveled by the centrosome. (B-C) Time lapse of a pre-abscising human (HeLa) cell (B) and zebrafish embryo cell (C). Inset with associated track (color gradient represents time) of distance traveled by centrosome (cell body centered for each time point) using equation in (A). Scale bar, 10 μm. (D) Western blot immunolabeled for Rab11 and pericentrin with GAPDH loading control in wildtype cells, Rab11-null cells and Rab11-null cells ectopically expressing mCherry-Rab11, - Rab11(S25N) and -Rab11(Q70L). (E-F) Representative images of fixed interphase human (HeLa) cells with single nuclei or bi-nuclei labeled using DAPI (gray, left; cyan, merge; E). Centrin-GFP (magenta) shown (E). Orange dashed lines indicate cell boundaries. Scale bar, 10 μm. Percentage of binucleate cells calculated across n=3 experiments for n>100 cells counted per experiment (F). One-way ANOVA with Dunnett’s multiple comparison to wildtype control cells shown, **p<0.01. (G) Total distance traveled by centrosome during pre-abscission measured across n>15 cells per condition. Scatter plot with the median (orange dashed line) and quartiles (dark lines) shown. Two-tailed student’s t-test, ****p<0.0001. (H-K) Rab11-null cells ectopically expressing mCherry- Rab11 and -Rab11(Q70L) shown (fire, LUT). ROI photobleached highlighted in inset (right), pre- and post-FRAP. Scale bar 10 μm (H). FRAP trace of mCherry-Rab11 and - Rab11(Q70L) shown for n>15 cells per condition (I). (J-K) mCherry-Rab11 and - Rab11(Q70L) mobile fraction (J) and half-life (t1/2, K) calculated and presented as violin plot with median (orange dashed line) and quartiles (dark lines) for n>15 cells per condition. Two-tailed student’s t-test, **p<0.01, ****p<0.001. Statistical results detailed in Table S2.

**Figure S3:**
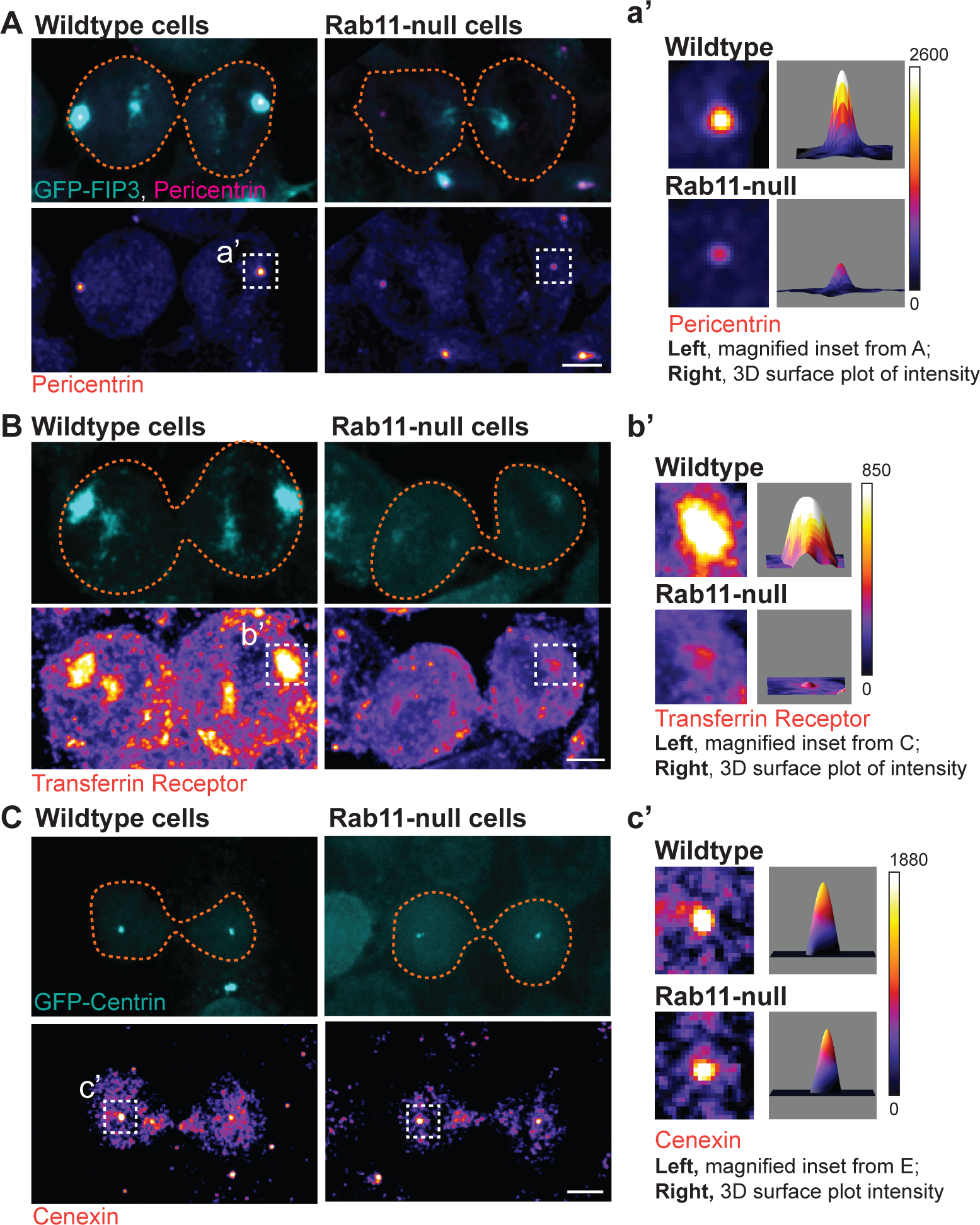
Rab11-GTP cycling is required for centrosome protein, pericentrin, centrosome localization. (A-C) Human (HeLa) pre-abscising cells expressing GFP-FIP3 (cyan, A, B) or centrin-GFP (cyan, C) were fixed and immunostained for pericentrin (fire LUT, A), transferrin receptor (fire LUT, B), and cenexin (fire LUT, C). Magnified insets (3x) shown on right (a’, b’, c’) with associated three-dimensional surface plot of intensity. Scale bar, 5 μm.

**Figure S4:**
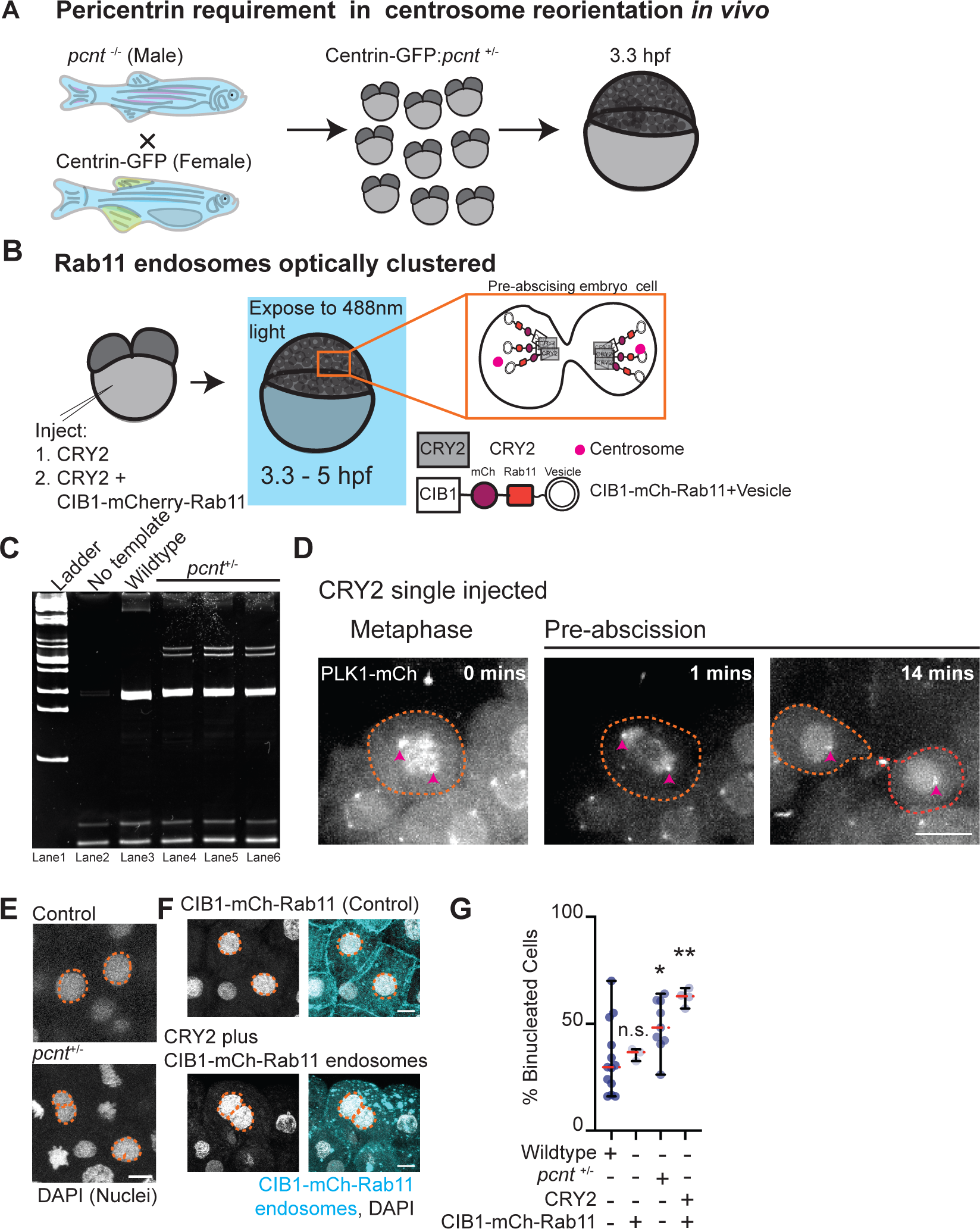
Pericentrin and Rab11 coordinate centrosome movement and number during mitotic exit *in vivo*. (A) Model depicting generation of *pcnt*^+/-^ embryos with labeled centrosomes (centrin-GFP) to test the role of pericentrin in centrosome reorientation and centrosome number *in vivo*. (B) Model depicting optogenetic clustering protocol for Rab11 endosomes. In short, embryos are injected with CRY2 and/or CIB1-Rab11 mRNAs, exposed to blue light 3.3-5 hpf causing a heterointeraction between CRY2 and CIB1, and imaged. Orange inset depicts cells imaged within the embryo. Pink circle, centrosome. (C) Ethidium bromide stained acrylamide gel electrophoresis showing *pcnt*^+/+^ and *pcnt*^+/-^ genotypes with ladder (lane 1), no template (lane 2), DNA extracted from wildtype zebrafish (*pcnt*^+/+^, lane 3) and *pcnt*^+/-^ embryos used in experimental analysis in Figure 4 (lane 4, 5, 6). (D) Time-lapse projections depicting a control dividing cell injected with CRY2 and PLK1-mCherry (gray). Pink arrow, centrosome. Dashed lines, cell boundaries. Scale bar, 10 μm. (E-F) Representative images of fixed interphase zebrafish embryonic cells with single nuclei or bi-nuclei labeled using DAPI in control, *pcnt*^+/-^, CIB1-mCherry-Rab11 injected, and CRY2 plus CIB1-mCherry-Rab11 injected embryos (gray, nuclei outlined with dashed orange line; E-F). Scale bar, 5 μm. Percentage of embryos with cells containing binucleated cells was calculated across n>30 cells in n>3 embryos (G). Statistical results detailed in Table S4.

Video S1: Mitotic centrosomes reorient towards the cytokinetic bridge during pre-abscission. Related to Figure 1.

Timelapse imaging of a dividing zebrafish embryo cell. Yellow arrow notes centrosome (centrin-GFP, gray; PLK1-mCherry, cyan). Video acquired over fourteen minutes at two-minute intervals. Scale bar, 10 μm.

Video S2: Mitotic centrosomes containing Rab11-endosomes reorient towards the cytokinetic bridge during pre-abscission. Related to Figure 1.

Timelapse imaging of a human (HeLa) cell centrosomes (DsRed-PACT, magenta) and REs (GFP-FIP3, cyan) depicted. Video acquired over two hours at one and a half minute intervals. Scale bar, 10 μm.

Video S3: Rab11 GTPase function and centrosome localization is required for centrosome bridge directed movement. Related to Figure 2.

Timelapse video of wildtype (left) and Rab11-null (right) human cells. Pink arrow notes centrosome (centrin-GFP, inverted grays). Video acquired over three hours at four-minute intervals. Scale bar, 10 μm.

Video S4: Rab11 GTPase function and centrosome localization is required for centrosome bridge directed movement. Related to Figure 2.

Timelapse video of Rab11-null human cells ectopically expressing plus mCherry-Rab11 (left), plus mCherry-Rab11(S25N, middle) and plus mCherry-Rab11(Q70L, right). Pink arrow notes centrosome (centrin-GFP, inverted grays). Video acquired over two hours at three to four-minute intervals. Scale bar, 10 μm.

**Table S1.** Detailed statistical analysis results reported in Figure 1 and Figure S1. Related to STAR methods.

| Figures | Y-Axis | Category | n (cell) | n (embryo) | n (experiment) | Statistical Test | Parameters | Result | P-value |
| --- | --- | --- | --- | --- | --- | --- | --- | --- | --- |
| 1C | Daughter cells with bridge directed centrosome movement (%) | Zebrafish embryo cell (Centrin-GFP) | n=26 | n=4 |  | N/A | N/A | N/A | N/A |
|  |  | Zebrafish embryo cell (PLK1-mCherry) | n=40 | n=4 |  |  |  |  |  |
|  |  | Human cell (HeLa) Centrin-GFP | n=15 | N/A | n=3 |  |  |  |  |
|  |  | Human cell (HeLa) Ds-Red-PACT | n=21 | N/A | n=4 |  |  |  |  |
| S1G | Fold change of photoconverted Dendra-Rab11 increase at the cytokinetic bridge | Before Photobleaching | n=6 | N/A | n=4 | Two-tailed Student's t-test | t=6.029, dF=10 | *** | 0.0001 |
|  |  | After Photobleaching |  | N/A |  |  |  |  |  |

**Table S2.** Detailed statistical analysis results reported in Figure 2 and Figure S2. Related to STAR methods

| Figures | Y-Axis | Category | n (cell) | n (experiment) | Statistical Test | Parameters | Result | p-value |
| --- | --- | --- | --- | --- | --- | --- | --- | --- |
| 2C | Daughter cells with bridge directed centrosome movement (%) | Wildtype | n=25 | n=5 | N/A | N/A | N/A |  |
|  |  | Rab11-null cells | n=35 | n=5 |  |  |  |  |
|  |  | Rab11-null cells, plus mCh-Rab11 | n=7 | n=3 |  |  |  |  |
|  |  | Rab11-null cells, plus mCh-Rab11(S25N) | n=9 | n=3 |  |  |  |  |
|  |  | Rab11-null cells, plus mCh-Rab11(Q70L) | n=6 | n=3 |  |  |  |  |
| 2F | Directional displacement of centrosome towards the cytokinetic bridge (µm) | Wildtype | n=9 | n=5 | One Way ANOVA | F (4, 87) = 6.21 |  |  |
|  |  | Rab11-null cells | n=15 | n=5 |  |  | **** | 0.0002 |
|  |  | Rab11-null cells, plus mCh-Rab11 | n=7 | n=3 |  |  | n.s. | 0.1019 |
|  |  | Rab11-null cells, plus mCh-Rab11(S25N) | n=9 | n=3 |  |  | **** | 0.0009 |
|  |  | Rab11-null cells, plus mCh-Rab11(Q70L) | n=6 | n=3 |  |  | **** | 0.0006 |
| Sup2F | % Binucleated cells | Wildtype | n>100 | n=3 | One Way ANOVA | F (2,6) = 18.07 |  |  |
|  |  | Rab11-null cells | n>100 | n=3 |  |  | ** | 0.0021 |
|  |  | Rab11-null cells, plus mCh-Rab11 | n>100 | n=3 |  |  | n.s. | 0.3245 |
| S2G | Total distance traveled by centrosome post anaphase exit (μm) | Wildtype | n=9 | n=4 | Two-tailed Student's t-test | t=5.000, df=32 | **** | <0.0001 |
|  |  | Rab11-null cells | n=8 | n=4 |  |  |  |  |
| S2J | Mobile Fraction (%) | Rab11-null cells, plus mCh-Rab11 | n=18 | n>3 | Two-tailed Student's t-test | t=3.208, df=30 | ** | 0.0032 |
|  |  | Rab11-null cells, plus mCh-Rab11(Q70L) | n=15 | n>3 |  |  |  |  |
| S2K | Half-life (t1/2, sec) | Rab11-null cells, plus mCh-Rab11 | n=18 | n>3 | Two-tailed Student's t-test | t=4.594, df=31 | **** | <0.0001 |
|  |  | Rab11-null cells, plus mCh-Rab11(Q70L) | n=14 | n>3 |  |  |  |  |

**Table S3.** Detailed statistical analysis results reported in Figure 3 and Figure S3. Related to STAR methods.

| Figures | Y-Axis | Category | n (centrosomes) | n (experiments) | Statistical Test | Parameters | Result | p-value |
| --- | --- | --- | --- | --- | --- | --- | --- | --- |
| 3D | Ratio of Rab11-null centrosome intensity / Wildtype intensity (A.U.) | Centrin | n>30 | n=3 | One Way ANOVA | F (4, 10) = 16.55 |  |  |
|  |  | Cenexin |  | n=3 |  |  | n.s. | 0.9991 |
|  |  | Pericentrin |  | n=3 |  |  | * | 0.0352 |
|  |  | GFP-FIP3 |  | n=3 |  |  | *** | 0.0008 |
|  |  | Transferrin Receptor |  | n=3 |  |  | *** | 0.0004 |
| 3F | Pericentrin normalized intensity pre-abscising cells (A.U.) | Wildtype | n=38 | n=5 | One Way ANOVA | F (4, 242) = 18.19 |  |  |
|  |  | Rab11-null cells | n=43 | n=5 |  |  | **** | <0.0001 |
|  |  | Rab11-null cells, plus mCh-Rab11 | n=62 | n=4 |  |  | n.s. | 0.6632 |
|  |  | Rab11-null cells, plus mCh-Rab11(S25N) | n=61 | n=4 |  |  | **** | <0.0001 |
|  |  | Rab11-null cells, plus mCh-Rab11(Q70L) | n=43 | n=3 |  |  | **** | <0.0001 |

**Table S4.**
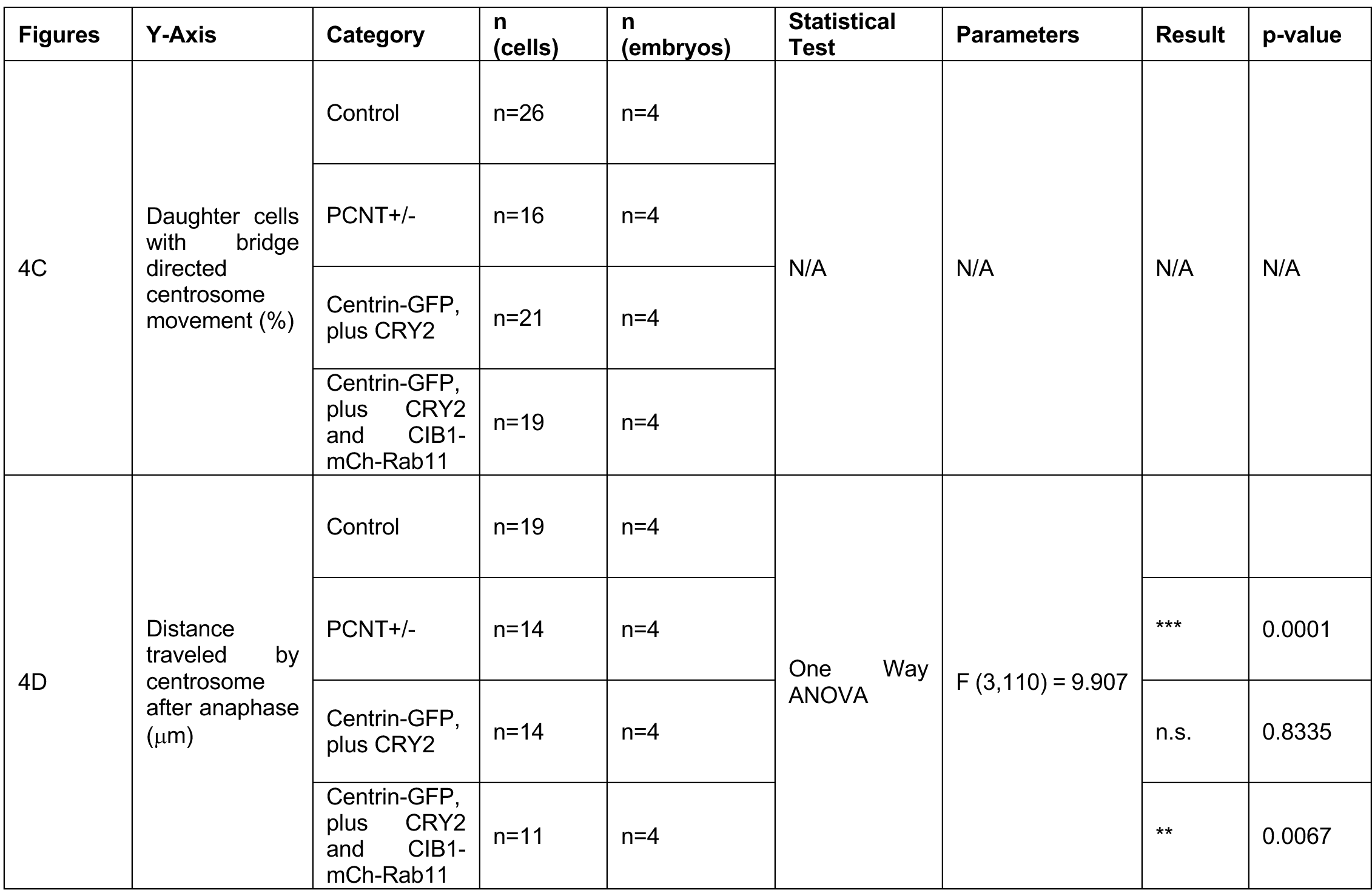
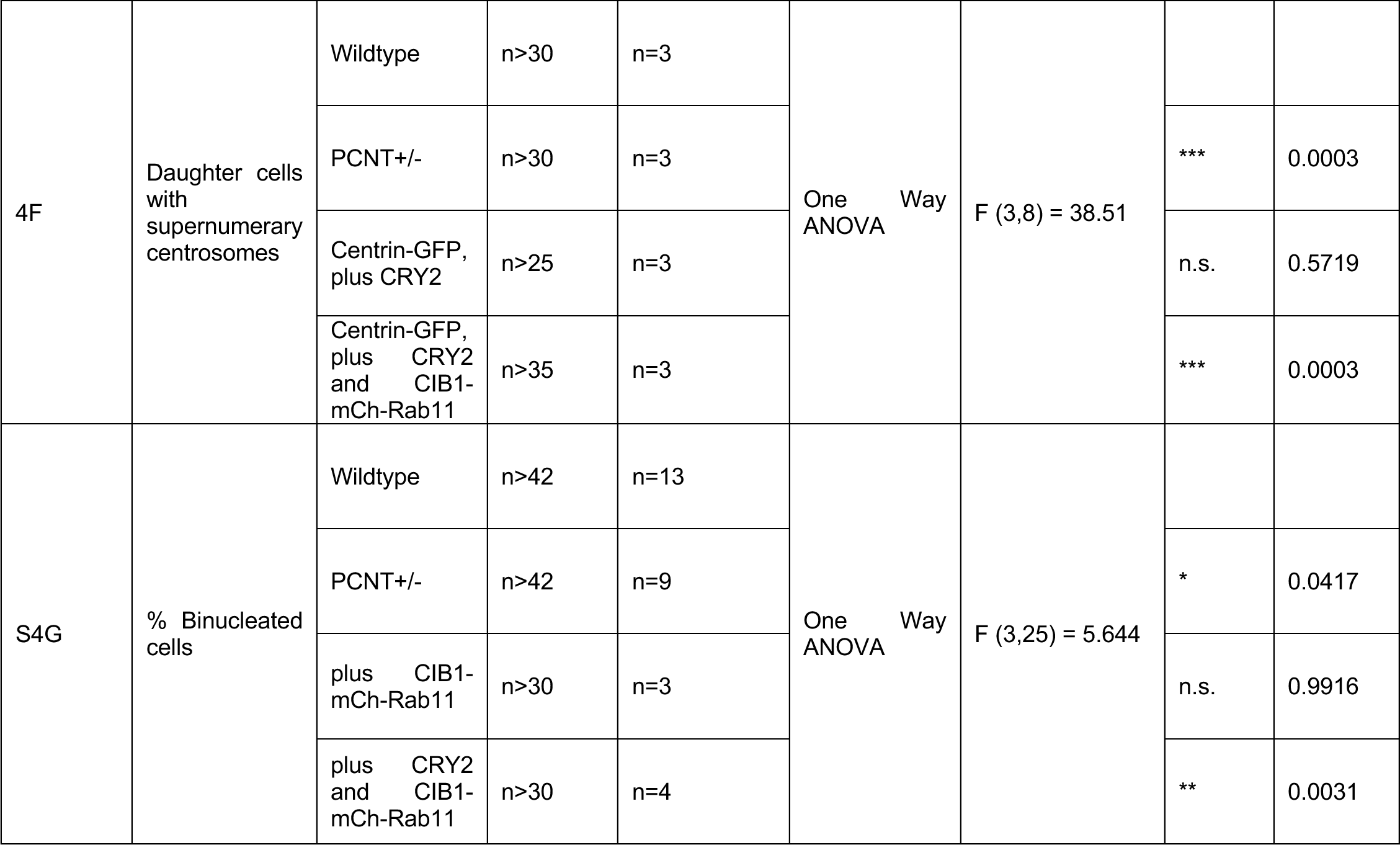
Detailed statistical analysis results reported in Figure 4 and Figure S4. Related to STAR methods.

| Figures | Y-Axis | Category | n (cells) | n (embryos) | Statistical Test | Parameters | Result | p-value |
| --- | --- | --- | --- | --- | --- | --- | --- | --- |
| 4C | Daughter cells with bridge directed centrosome movement (%) | Control | n=26 | n=4 | N/A | N/A | N/A | N/A |
|  |  | PCNT+/- | n=16 | n=4 |  |  |  |  |
|  |  | Centrin-GFP, plus CRY2 | n=21 | n=4 |  |  |  |  |
|  |  | Centrin-GFP, plus CRY2 and CIB1-mCh-Rab11 | n=19 | n=4 |  |  |  |  |
| 4D | Distance traveled by centrosome after anaphase (μm) | Control | n=19 | n=4 | One Way ANOVA | F (3,110) = 9.907 |  |  |
|  |  | PCNT+/- | n=14 | n=4 |  |  | *** | 0.0001 |
|  |  | Centrin-GFP, plus CRY2 | n=14 | n=4 |  |  | n.s. | 0.8335 |
|  |  | Centrin-GFP, plus CRY2 and CIB1-mCh-Rab11 | n=11 | n=4 |  |  | ** | 0.0067 |
| 4F | Daughter cells with supernumerary centrosomes | Wildtype | n>30 | n=3 | One Way ANOVA | F (3,8) = 38.51 |  |  |
|  |  | PCNT+/- | n>30 | n=3 |  |  | *** | 0.0003 |
|  |  | Centrin-GFP, plus CRY2 | n>25 | n=3 |  |  | n.s. | 0.5719 |
|  |  | Centrin-GFP, plus CRY2 and CIB1-mCh-Rab11 | n>35 | n=3 |  |  | *** | 0.0003 |
| S4G | % Binucleated cells | Wildtype | n>42 | n=13 | One Way ANOVA | F (3,25) = 5.644 |  |  |
|  |  | PCNT+/- | n>42 | n=9 |  |  | * | 0.0417 |
|  |  | plus CIB1-mCh-Rab11 | n>30 | n=3 |  |  | n.s. | 0.9916 |
|  |  | plus CRY2 and CIB1-mCh-Rab11 | n>30 | n=4 |  |  | ** | 0.0031 |

